# Sex differences in alpha galactosidase protein processing and its impact on disease severity in Fabry disease

**DOI:** 10.1101/2025.09.16.676554

**Authors:** L Lavalle, H Kurdi, D Moreno Martinez, S Mangrati, DA Hughes

## Abstract

Fabry disease (FD) is an X-linked disorder due to mutations in the α-galactosidase A (GLA) gene. The condition is characterized by low GLA activity and accumulation of toxic sphingolipids. Some patients present full disease symptoms whereas others have one system affected, generally the heart or kidney. This suggests that a mutation in the GLA gene is necessary to cause FD, but other factors may contribute to its clinical expression.

To investigates the impact of GLA mutant protein processing on GLA activity and disease expression, 26 individuals with FD (14 males) were studied. Clinical outcomes explored included the Mainz Severity Score Index (MSSI), the Age-Adjusting Severity Scores (AASS), glomerular filtration rate (GFR), and left ventricular mass index (LVMI). Globotriaosylsphingosine (lyso-Gb3) data was available for a subset of participants.

Whole cell lysates were employed to assess GLA activity and GLA protein levels. Additionally, endoglycosidase H digestion was performed on cell lysates to quantify each GLA protein form: immature 50 kDa endoplasmic reticulum form and mature 46 kDa lysosomal form. The latter was employed to calculate lysosomal GLA activity and its relationship with clinical outcomes was studied.

Fabry participants exhibited more of the immature form of GLA protein (0.23 vs 0.66, p= 0.04). Female patients exhibited higher total GLA activity (19.0 vs 5.1 nmol/hr/mg), total GLA protein levels 0.13 vs 0.50 GLA/LC, p= 0.003), and 46 kDa mature lysosomal form levels than male patients (0.53 vs 0.10, p= 0.001). Additionally, females showed a significant correlation between the GLA mature form and GLA activity (r2= 0.59, p= 0.04). Consistently, lysosomal GLA activity exhibited significant associations with MSSI (r2= -0.66, p= 0.02) and GFR (r2= 0.59, p= 0.04) only in this sex group.

These results suggest that total GLA protein levels are linked to the severity of FD manifestations, particularly in females via enzyme activity.

## 1.0 Introduction

Fabry disease (FD) is a rare genetic disorder characterized by low α-galactosidase A (GLA) activity and high variability in disease expression. While some patients exhibit full disease symptoms (’classical’ FD) others may only experience involvement of a single organ system, typically affecting the heart or kidneys (‘non-classical’ or ‘later onset’ FD) (Hughes, 2017). Predicting the disease course based on genotype presents challenges as some missense GLA mutations can give rise to a range of phenotypes, even when associated with high preserved GLA activity (van der Tol et al., 2014). The deficit of GLA activity results in the accumulation of globotriaosylceramide (Gb3) and its derivate globotriaosylsphingosine (lyso-Gb3). These are toxic sphingolipids and while Gb3 is considered a hallmark of FD, higher levels of the latter correlate to increased clinical severity and disease progression (Smid et al., 2015).

Patients with FD can present with a wide range of signs and symptoms. Males with the classical form of the disease will have very low (<1%) or absent GLA activity and therefore can be diagnosed by enzyme activity analysis (Reuser et al., 2011). For heterozygous females, GLA activity can remain within a normal range for which genotyping and accurate interpretation of GLA variant is required. The gold standard method to measure GLA activity employs total protein extracts from peripheral blood leukocytes, whole blood, dried blood spots, or plasma (Desnick RJ, 2007; Gal, Hughes, & Winchester, 2011; Hughes, 2017; Massaccesi et al., 2011).

GLA protein is synthesized and translocated into the endoplasmic reticulum (ER) lumen for glycosylation and folding (Lemansky, Bishop, Desnick, Hasilik, & von Figura, 1987). The glycan serves as a folding barcode for the ER quality control, allowing recognition of the folding status of the polypeptide. Based on this, local chaperons will continue folding the protein or tag it for degradation. When this process is compromised, it leads to accumulation of misfolded protein within the ER lumen (Shenkman M, 2019). Using a Drosophila melanogaster models of FD, the Horowitz group expressed the classical variants A156V and A285D displaying 4.3% in vitro GLA activity and no detectable activity, respectively. While most of the A285D protein was retained in the ER, a significant fraction of the A156V protein was able to exit this organelle (Braun et al., 2019; Braunstein et al., 2020).

Currently, GLA activity is the most accurate predictor of clinical severity of FD; however, the minimal activity required to avoid substrate buildup and preserve lysosomal homeostasis is unknown (Lukas et al., 2013; Schiffmann, Fuller, Clarke, & Aerts, 2016). It has been suggested that lysosomal hydrolases possess a significant functional reserve, with substrate storage only occurring when GLA activity falls below a critical threshold of 10% - 15% of normal levels (Leinekugel, Michel, Conzelmann, & Sandhoff, 1992). Nevertheless, recent work has raised questions about traditional methods to measure lysosomal enzymes’ activities. This suggest that the standard in vitro assay measures aggregate GLA activity rather than the activity within the lysosome, leading to an overestimation of the enzyme activity available to patients (Cecioni et al., 2022).

This study investigates the relation between GLA protein variants and their intracellular processing to understand whether variation in these pathways impact GLA activity and clinical outcomes, including kidney function, left ventricular mass index (LVMI), overall disease severity by the Mainz severity score index (MSSI), and its age and gender corrected counterpart, the age adjusting severity score (AASS). To this end, we cultured fibroblast cell lines and PBMCs from individuals with FD and measured total GLA activity and protein. Whole cell lysates were subjected to endo-H digestion to quantify each GLA protein form: immature 50 kDa ER form and mature 46 kDa lysosomal form. The latter was employed to calculate lysosomal GLA activity and its relationship with clinical outcomes was studied.

## 2.0 Methods

The study received ethical approval from the Health Research Authority (HRA) and Health and Care Research Wales (HCRW) (REC number: 20/WM/0329). Patients are required to give written consent for study participation. Study documents and ethical approval can be found in Appendix 1.

### 2.1 Clinical data collection and analysis

All participants with FD included in this study were documented to have at least one GLA mutation and had measurement of either plasma or leukocyte enzyme activity available on patients notes, done prior to initiation of treatment. Medical history of all patients were retrospectively assessed and, when available, clinical data prior to initiation of treatment was collected and employed to calculate clinical severity scores, i.e., MSSI and AASS, avoiding any influence of treatment effect on organ manifestations. Data regarding clinical events after treatment commencement were used to cluster individuals with FD into clinical signatures.

Cardiac data included electrocardiogram (ECG) and cardiac magnetic resonance (CMR). Left ventricular mass was calculated using the Devereux formula (Devereux et al., 1986) and LVMI was calculated and adjusted for height. Left ventricular hypertrophy (LVH) was define as an LVMI >/= 78 and 74 g/m^2^ in males and females, respectively (Chuang et al., 2014). Renal function was assessed by estimated glomerular filtration rate (GFR), calculated using the CKD Epidemiology Collaboration equation (CKD-EPI)(Lamb, Levey, & Stevens, 2013) and CKD was staged according to the Renal Association, UK. The presence of WMLs was evaluated by cerebral magnetic resonance imaging (MRI). Plasma lyso-Gb3 measurements were available for a subset of patients prior to treatment commencement.

#### Classical vs nonclassical FD

participants were classified as classical or late onset / nonclassical FD based on residual enzyme activity (only for males) and the presence or absence of characteristics Fabry symptoms, i.e., acroparesthesia, angiokeratoma, and / or cornea verticillate. Males were classified as having the classical phenotype if (1) their GLA activity was lower than 5% of the mean reference range and (2) had at least one of the characteristic Fabry symptoms. Alternatively, they were classified as nonclassical / late onset FD. For females, the presence of at least 1 characteristic Fabry symptom was sufficient to be consider as having a classical phenotype. Conversely, they were classified as nonclassical / late onset FD (Smid et al., 2014).

#### Clinical signatures

individuals with FD were subsequently clustered according to the predominant organ system affected, defining these as “clinical signatures”.

#### Fabry cardiomyopathy

was defined as (1) the presence of ECG abnormalities included Sokolow-Lyon voltage criteria for LVH, conduction abnormalities (long QT, short PR, and conduction blocks), repolarization abnormalities (T-wave inversions in at least two contiguous), increase voltages and arrhythmias including multifocal ventricular ectopics, ventricular fibrillation, non-sustained ventricular tachycardia, ventricular tachycardia, (2) LVH documented on image scan, and / or (3) the development of serious cardiac events included atrial fibrillation and other clinically significant arrythmia, implantation of an implantable cardiac defibrillator or pacemaker, and myocardial infarction (Nordin et al., 2019).

#### Fabry nephropathy

Latest expert reviews report characteristic FD features in kidney biopsies of Fabry patients with normal GFR and absence of proteinuria. These features include segmental podocyte foot processes effacement, focal sclerosis, Gb3 depositions, and vascular changes (Ortiz et al., 2008; Schiffmann et al., 2017). A more recent study reported glomerular hyperfiltration as a common feature in young patients with FD and highlighted its contribution to the loss of renal function and onset of nephropathy in type 2 diabetes, sickle cell anaemia, and in autosomal dominant polycystic kidney disease (Riccio et al., 2019). Therefore, in this study we defined Fabry nephropathy as (1) the presence of glomerular hyperfiltration (GFR> 130ml/min/1.73m2), (2) a low GFR: for adults under the age of 60 this was defined as an GFR< 90ml/min/1.73m2. For individuals aged 60 or older this was defined according to their age (Fernandes, Fernandes, Magacho, & Bastos, 2015), (3) proteinuria (>150mg/24hs) or an increased albumin creatinine ratio (> 3mg/mmol), and / or (4) the development of serious renal events including CKD stage 3A (GFR<59ml/min/1.73m2), dialysis, and kidney transplant.

#### Fabry cerebrovascular diseas

was define as (1) the presence of WML and (2) history of stroke or TIA.

### 2.2 Cell cultures

Cells were incubated in a humidified atmosphere with 5% CO2 at 37°C (Haraeus Instruments, Michigan, US).

Fibroblast cell lines GM00302, GM00881, GM00882 (Coriell Institute) were grown in Eagle’s Minimum Essential Medium with Earle’s salts and non-essential amino acids (EMEM; Stemcell Technologies; 36550), supplemented with 15% heat-inactivated FBS (Thermo Fisher; 26140079), and 1% Penicillin-Streptomycin (Sigma; P0781).

Peripheral blood mononuclear cells (PBMC) were isolated from 10 ml of blood by Lymphoprep™ (Stemcell Technologies; 07801) gradient separation. PBMCs were grown in Roswell Park Memorial Institute (RPMI) 1640 medium (Thermo Fisher; A10491-01) with 10% heat inactivated FBS. PBMCs for all FD patients and healthy controls (HC) were cultured for a total of 6 days with 2 media renewals.

### 2.3 Cell lysate preparation

Fibroblasts were harvested with TrypLE™ Express Enzyme (Life Technologies; 12604013) and transferred into centrifuge tubes. PBMCs were transferred into centrifuge tubes and spin down at 1500 rpm for 10 minutes at 4°C. Whole cell lysates were obtained by washing twice with Hanksʹ Balanced Salt solution (HBSS; Sigma; H8264). Then, 100 ul of mammalian cell lysis solution (GE Healthcare; 28-941279) was added, supplemented with 1% protease cocktail inhibitor (Sigma; P8340) and 1% of 100mM phenylmethylsulfonyl fluoride (17.4mg/ml in DMSO; Sigma). Total protein quantification was determined using a bicinchoninic acid protein assay kit (BCA; Sigma; BCA1) and supernatants were stored at -80°C until analysis.

### 2.4 GLA activity assay

GLA activity was measured in whole cell lysates samples containing 2 ug of total protein by fluorometric technique (Kizhner et al., 2015). 4-methylumbelliferyl-α-galactopyranoside (Sigma; M7633) was used as substrate and N-acetyl-D-galactosamine (Sigma; A2795) as an inhibitor of α-D-galactosidase B in 0.15 M McIlvaine’s buffer (pH 4.7). The reaction was terminated by adding 1.0 M glycine solution (pH 10.5).

### 2.5 Western blot analysis

Lysates containing 10 ug of total protein were combined with 10 ul 4X Bolt™ LDS Sample Buffer (Invitrogen™; B0007), 4 ul of 10X Bolt™ Sample Reducing Agent (Invitrogen™; B0009), 6 ul distilled H20 and gently mixed. Samples were then incubated at 90°C for 5 minutes and centrifuged at 13000 rpm for 30 seconds. Samples were loaded on a Bolt™ 4-12% Bis-Tris Plus Gels (Invitrogen™; NW04120BOX) and the gel was run at 200 volts for 32 minutes. 10 ul of protein ladder (SeeBlue™ Plus2 Pre-stained Protein Standard; Invitrogen™; LC5925) was employed. Proteins were transferred to a nitrocellulose blotting membrane (GE Healthcare Life Sciences; 10600002) and this was blocked using 10 ml of blocking buffer (5% milk solution in 0.1% Tween 20 PBS (Sigma; 70166)) at room temperature for an hour. Then the membrane was treated with primary antibodies labelled with horseradish peroxidase: anti-GLA monoclonal antibody (1.3 ng/ul; Bio-techne, NBP2-90048H) and anti-sodium potassium ATPase antibody (0.1 ng/ul; Abcam, ab185065) as loading control (LC). Development of the membrane was done by pipetting 2 ml of enhanced chemiluminescent substrate solution (SuperSignal™ West Pico PLUS Chemiluminescent Substrate; Thermo Fisher, 34577) at room temperature. Bands were visualised on a BioRad Molecular Imager (Invitrogen™) and signal quantifications were done in pre-saturating blots.

### 2.6 Endoglycosidase H Sensitivity

According to manufacturer’s instructions, samples of cell lysate containing 20 ug of total protein were subjected to an overnight incubation with endoglycosidase H (endo H; P0702L, New England BioLabs. In parallel, undigested control samples were prepared by using water instead of the enzyme and these were incubated alongside the experimental samples. Samples were stored at -80°C until western blot analysis. The assay was done as described above, using 10 ug of total protein per lane, and the intensity of the entire GLA band was quantified. By comparing the undigested control sample (in endo-H negative lane), the endo-H sensitive band was identified (in the endo-H positive lane), and its signal was measured. Endo-H resistant fraction was calculated as shown below, in the endo-H positive lane (see fig. 8A and fig. 9A).

### 2.7 Statistical analysis

All data was analysed using GraphPad Prism 8.0™ (GraphPad Software Inc., California, USA). Parametric data is presented as the mean and standard deviation, while non-parametric data is represented using the median and interquartile range, with scatter plots visualizing individual data points. N value is defined as the number of repeats for each patient and for each independent cell culture experiment. Difference between two discrete populations were analysed by Mann-Whitney U tests and difference between three or more datasets was analysed by Kruskal-Wallis test with post hoc Dunn’s test. Relationship between two continuous variables was determined using Spearman’s rank correlation. P<0.05 was considered to be significant.

## 3.0 Results

### 3.3.1 Total GLA protein and activity in fibroblasts from controls and subjects with FD

GLA activity was reduced in both Fabry derived fibroblasts but to a different extent, while R301G activity was reduced to 18.1% control activity, R220X displayed only 2.6%. To further characterize these cell lines, β-Galactosidase (B-GAL) and Glucocerebrosidase (GBA) activities were also measured. The former showed to be reduced in fibroblasts derived from Fabry subjects, with the R220X and R301G displaying 80.2% and 89.3% control activity, respectively. GBA activity was also measured and only showed a decrease in activity in the R220X cell line, to 90.2% control activity (figure 1B).

**Figure 1.**
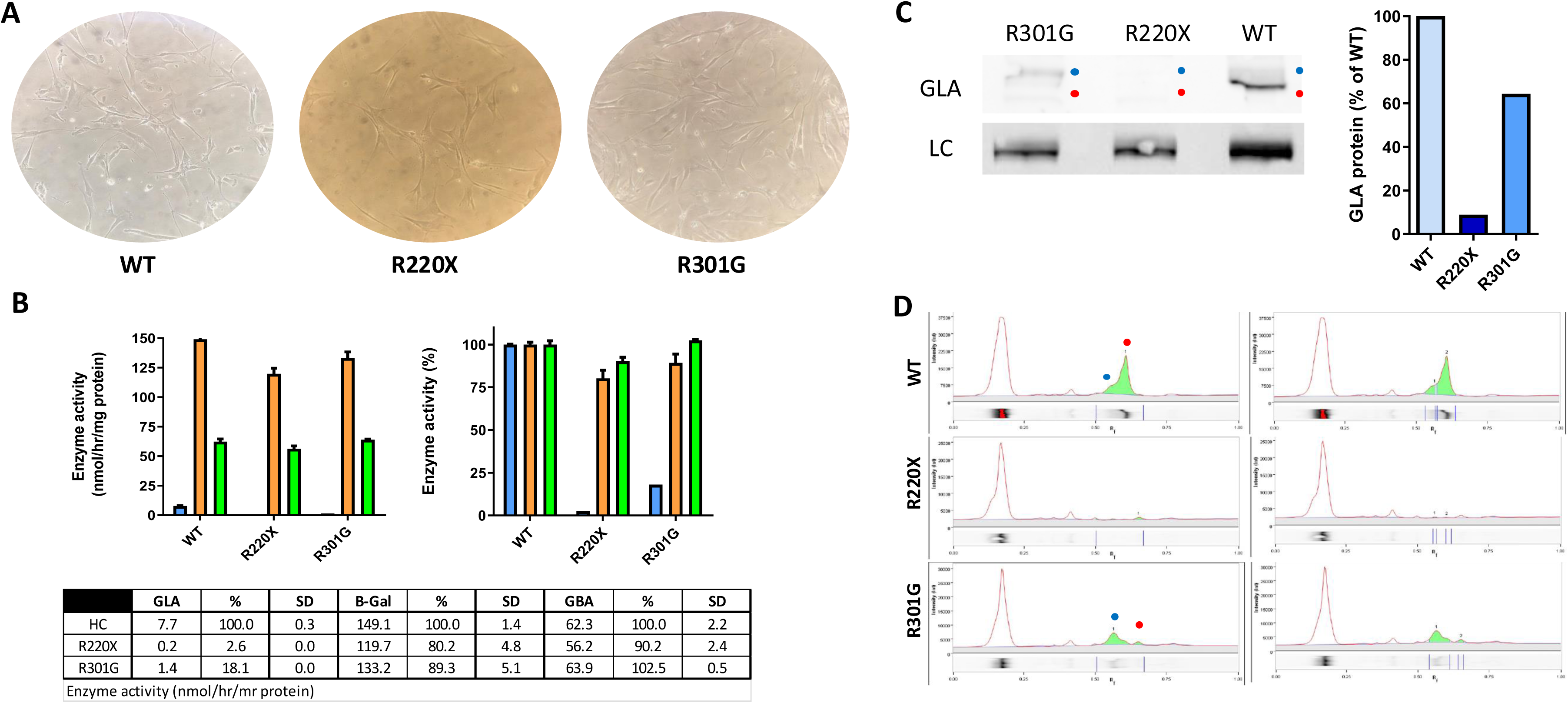
α-galactosidase A expression in control and FD fibroblasts. **(A)** Micrographs of fibroblast derived from 2 subjects with FD: Arg220Ter (R220X) and Arg301Gly (R301G) and an apparently healthy individual with a wild type (WT) GLA variant. Images were taken at 20X magnification at approximately 40% confluence in a T75 flask. **(B)** Samples containing 20 ug of protein were tested for total α-galactosidase A (GLA), β-Galactosidase (B-Gal), and Glucocerebrosidase (GBA) activities (n= 3). **(C)** Lysates containing 30 ug of protein were subject to SDS-PAGE and interaction with anti-GLA antibody and anti-sodium potassium ATPase antibody, as a loading control (LC). The blue dot indicated the higher molecular weight fraction (50 kDa ER retained) of the protein and the red dot indicates the lower molecular wight 46 kDa lysosomal fraction. To quantify the results, GLA protein intensity in each lane was divided by that of LC in the same lane, and the number obtained for WT was

Western blot analysis of lysates showed lower levels of GLA protein in Fabry derived fibroblasts in comparison to control. Moreover, the level of the nonsense mutant was lower than that of R301G variant (Figure 1C). This assay evidenced the appearance of a minor band (red dot) in the control sample with a major upper band (blue dot), two bands in the R301G lysate, and only one small band in the R220X mutant (1C and D).

#### 3.3.2 Total GLA protein and activity in controls and subjects with FD

The patient cohort comprised 27 Fabry disease patients (14 males and 12 females) harbouring a total of 16 different mutations (table 1A and B).

**Table 1A.**
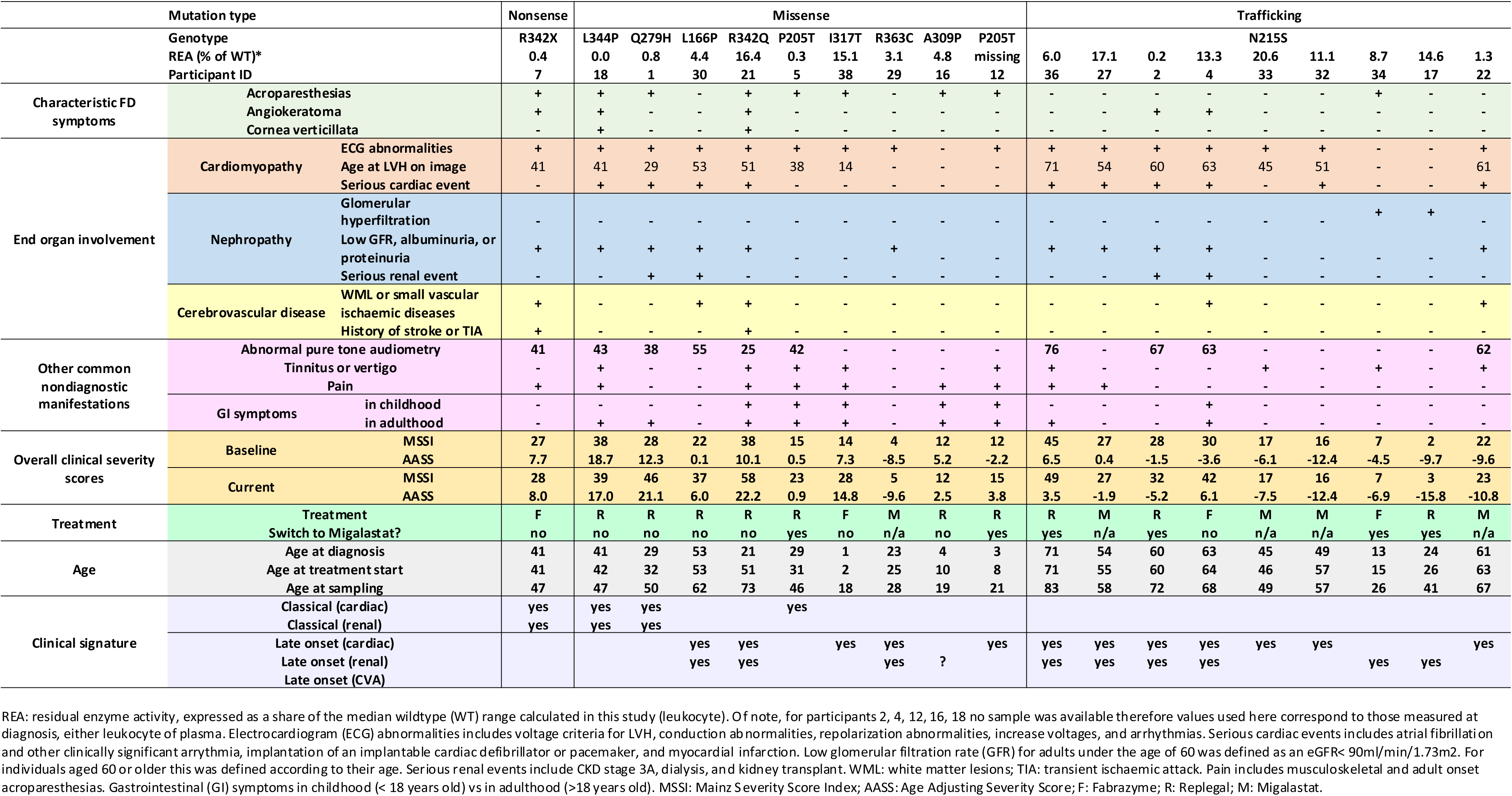
Males.

**Table 1B.**
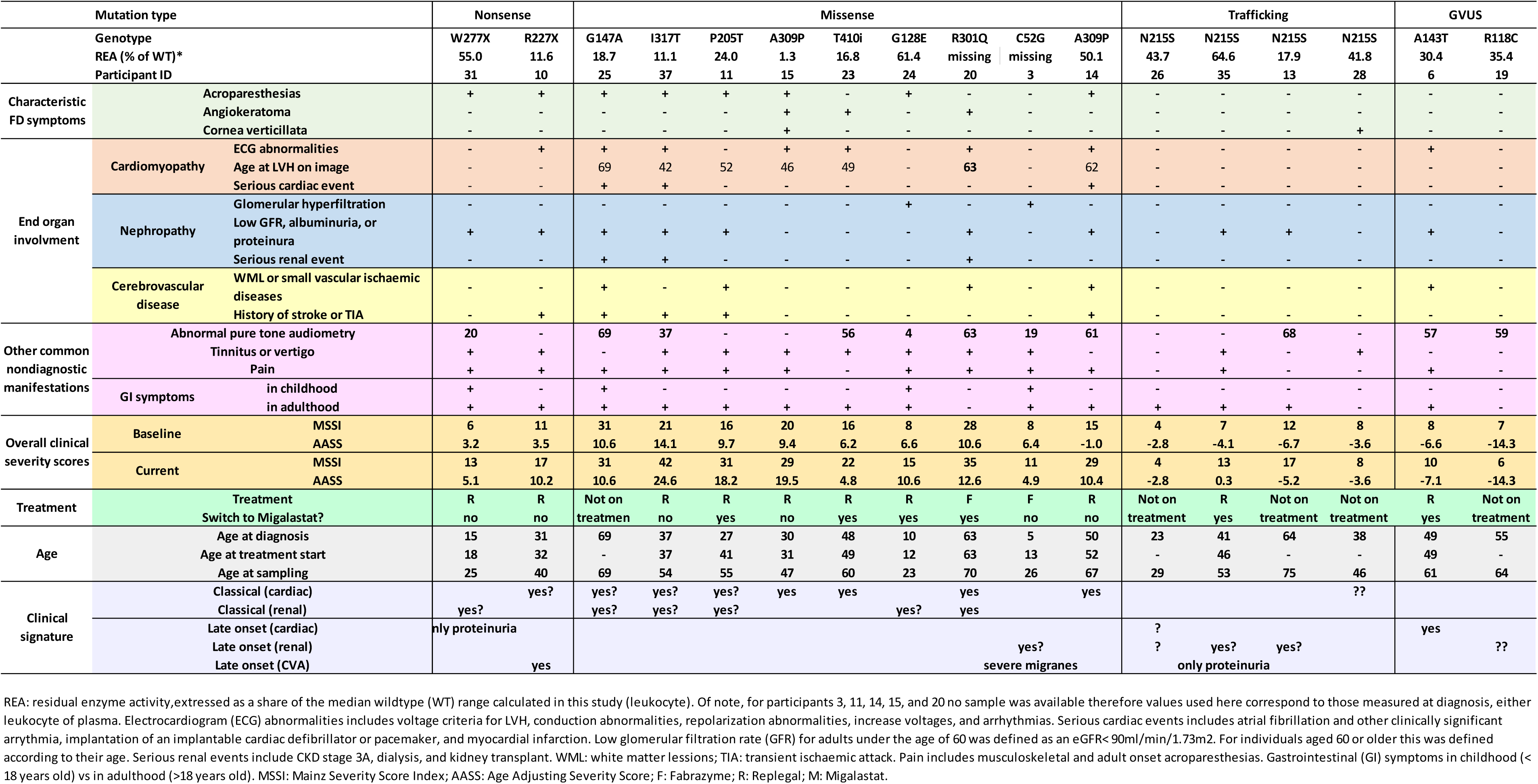
Females.

To understand the impact of culture on GLA activity and protein, PBMCs from HCs (1 female and 3 males) were isolated. Half were cultured as explained in the methods section, while the other half was immediately processed for protein extraction (following the same procedure used for cultured cells, as explained in the methods section). The median GLA activity following culture was 74.2 nmol/hr/mg (796.5% of normal activity in uncultured cells, n= 4, p< 0.03, fig 2A).

This increase in GLA activity following 6 days culture was consistent with the increased intensity of GLA protein band by western blot (715.9% vs 100%, n=4, p= 0.03, fig 2B). In all HC cell extracts, two forms of GLA protein were identified, a broad band ranging from 50.7 to 44.9 kDa (average 48.3 kDa, corresponding to the ∼50 kDa precursor form of GLA protein, ‘Upper’ band, fig 2C) and a smaller band ranging from 43.1 to 47 kDa (average 45.8 kDa, representing the ∼46 kDa mature lysosomal form, ‘Lower’ band, fig 2C), as previously reported (Hamanaka et al., 2008; LeDonne, Fairley, & Sweeley, 1983; Lemansky et al., 1987; Shin et al., 2008). While the latter represented most of the GLA protein band in uncultured cells (65.3% vs 33.4% n= 4, p< 0.03), this shifted after 6 days culture, with the 50 kDa immature form accounting for the majority form (99.6% vs 0.41% n= 4, p< 0.03, fig 2C). To determine if the 50 kDa precursor was active, the individual GLA protein forms were quantified relative to the loading control of the lane and plotted against corresponding GLA activity values. This showed that after culture higher amount of the precursor corresponded with higher levels of enzyme activity, suggesting that this form is enzymatically active, at least in the in vitro GLA activity assay employed in this study (fig. 2D).

**Fig. 2.**
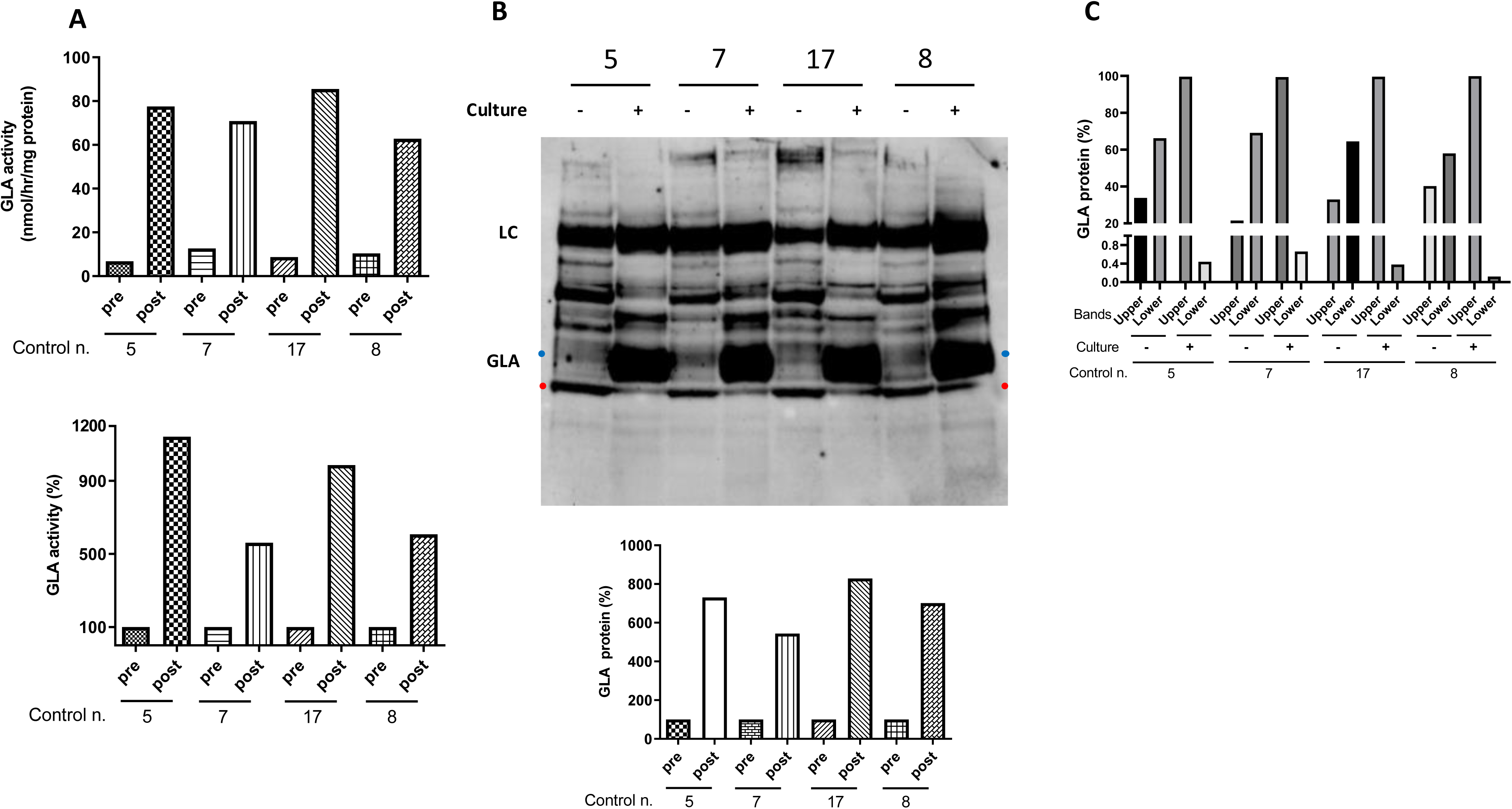
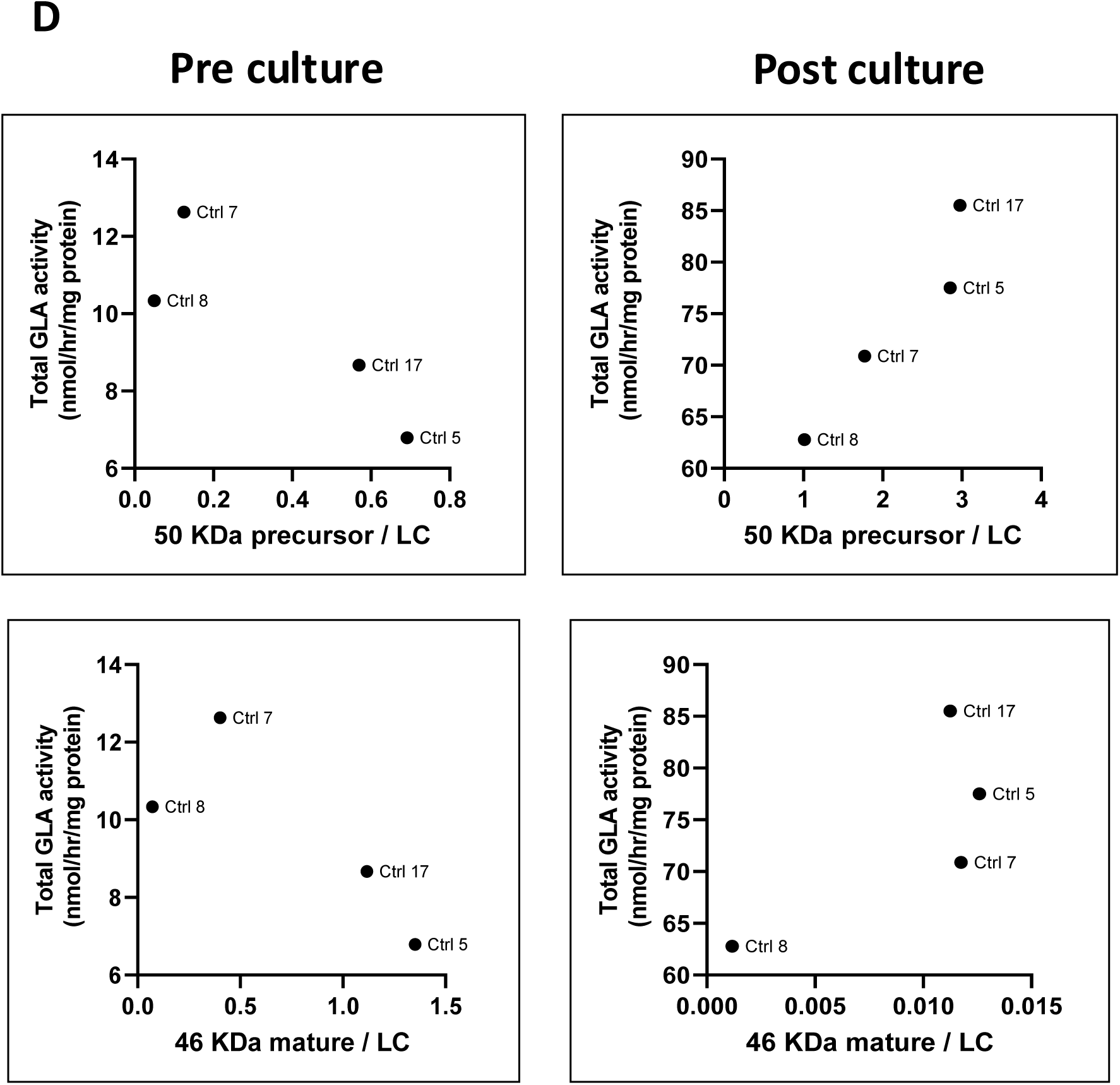
Impact of culture on α-galactosidase A activity and protein level. Peripheral blood mononuclear cells (PBMCs) were isolated from 4 healthy controls (HC), a female (HC 5) and 3 males (HC 7, 8, 17). For each participant, half of the cells obtained were cultures for 6 days with two media changes, and the other half was stored at -80 C until protein extraction. Cell lysates were prepared from both conditions for each control, i.e., pre and post culture. **(A)** Volumes containing 2 ug of protein were tested for GLA enzyme activity as described in the methods section (n= 3). **(B)** Volumes containing 10 ug of protein were subject to SDS-PAGE and interaction with anti - α galactosidase A antibody (GLA, 1.3 ng/ul, Bio-Techne) and anti-sodium potassium ATPase antibody (0.1 ng/ul, Abcam) as a loading control (LC). For total GLA quantification, band intensity in each lane was divided by that of the LC in the same lane, and the number obtained for the pre culture condition was considered to be 100 (n= 1). **(C)** In order to quantify the two forms of GLA (∼50 kDa ‘Upper’, blue dot, and ∼46 kDa ‘Lower’, red dot) the intensity of each was measured and divided by the intensity of the entire GLA band in the same lane. **(D)** To determine whether the 50 kDa precursor endoplasmic reticulum (ER) form is active, each band was quantified relative to the LC in the same lane and plotted against the corresponding enzyme activity i.e., pre or post culture.

GLA activity was significantly higher in controls than in FD patients regardless of gender (males: 70.5 vs. 5.1 nmol/hr/mg protein, p< 0.0001 and females: 57.1 vs. 19.2 nmol/hr/mg protein, p= 0.02, fig. 3A), and female patients’ activity was higher than that of male patients’ (19.2 vs 5.1 nmol/hr/mg protein, p= 0.0007, fig. 3A). GLA protein was also higher in this patient group 0.5 vs 0.1 GLA/LC, p= 0.003, fig. 4D) but these females’ GLA protein level had no differences with their HC counterpart (fig. 4C). Conversely, male HC did show significantly higher GLA protein levels than male individuals with FD (1.8 vs 0.1 GLA/LC, p= 0.003, fig. 4B). While neither activity nor total GLA protein showed significant differences between genotypes, marked variation on total GLA protein amounts was noted in patient with FD of both sex groups, and even among subjects with the same genotype i.e., N215S (fig. 4C).

**Fig. 3.**
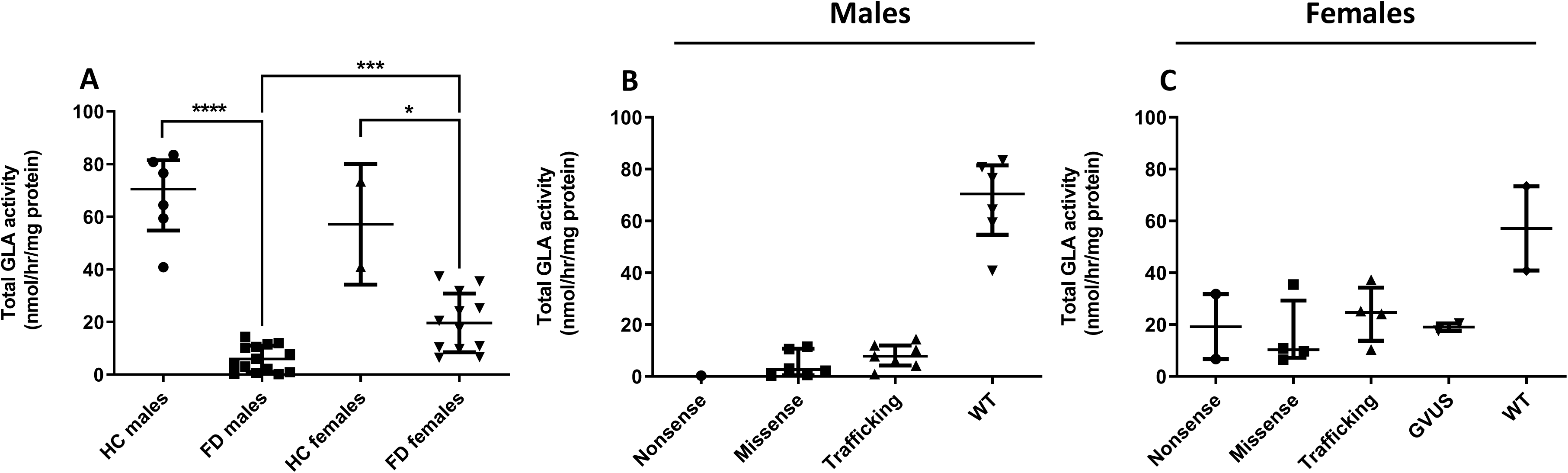
α-galactosidase A activity in mononuclear cells from control and participants with Fabry disease. **(A)** Cell lysates were prepared from peripheral blood mononuclear cells (PBMCs) of patients with Fabry disease (FD; 14 males and 13 females) and from healthy controls (HC; 6 males and 2 females). Samples containing 2 ug of total protein were tested for α-galactosidase A (GLA) activity as described in the methods section. Each point is the median of three wells and data presented in median and interquartile range per genotype and gender: **(B)** males and **(C)** females. Mutations include nonsense (R227X), trafficking (N215S), and genetic variants of unknown significance (GVUS; A143T and R118C). ****= p< 0.0001, *= p< 0.05.

**Fig. 4.**
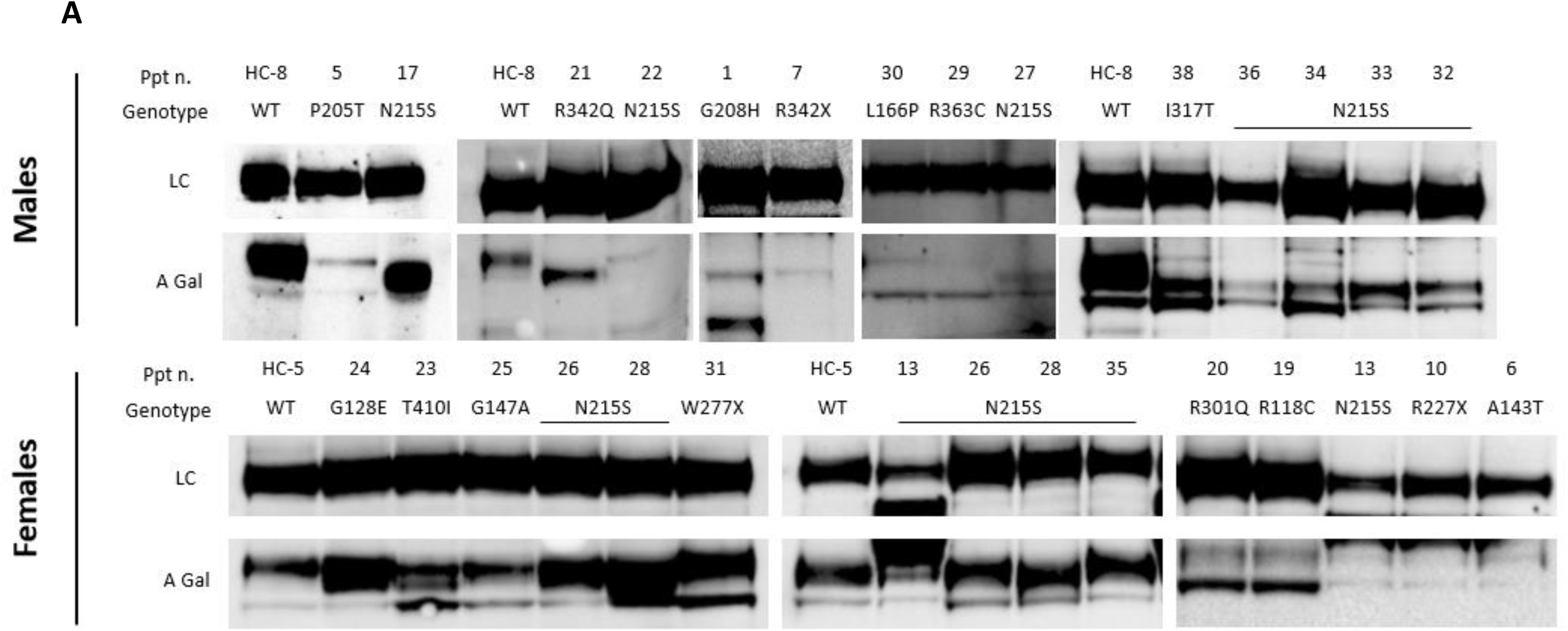
Quantification of α-galactosidase A in mononuclear cells from control and participants with Fabry disease by western blotting analysis. **(A)** Cell lysates containing 10 ug of protein prepared from peripheral blood mononuclear cells (PBMCs) of patients with FD and from healthy controls (HC) were subject to SDS-PAGE and interaction with anti - α-galactosidase A (GLA; 1.3 ng/ul, Bio-techne) antibody and anti-sodium potassium ATPase antibody (0.1 ng/ul, Abcam) antibody, as a loading control (LC). **(B)** For total GLA protein quantification, GLA band intensity in each lane was divided by that of LC in the same lane. **(C)** The median obtained for HCs was set as 100%. Data presented in median and interquartile range; n= 1 to 2 per FD patient, HC= 3 males (n=1, n=1, n=4) and 1 female (n=3). **= p< 0.01, ns= not significant.

**Fig. 4.**
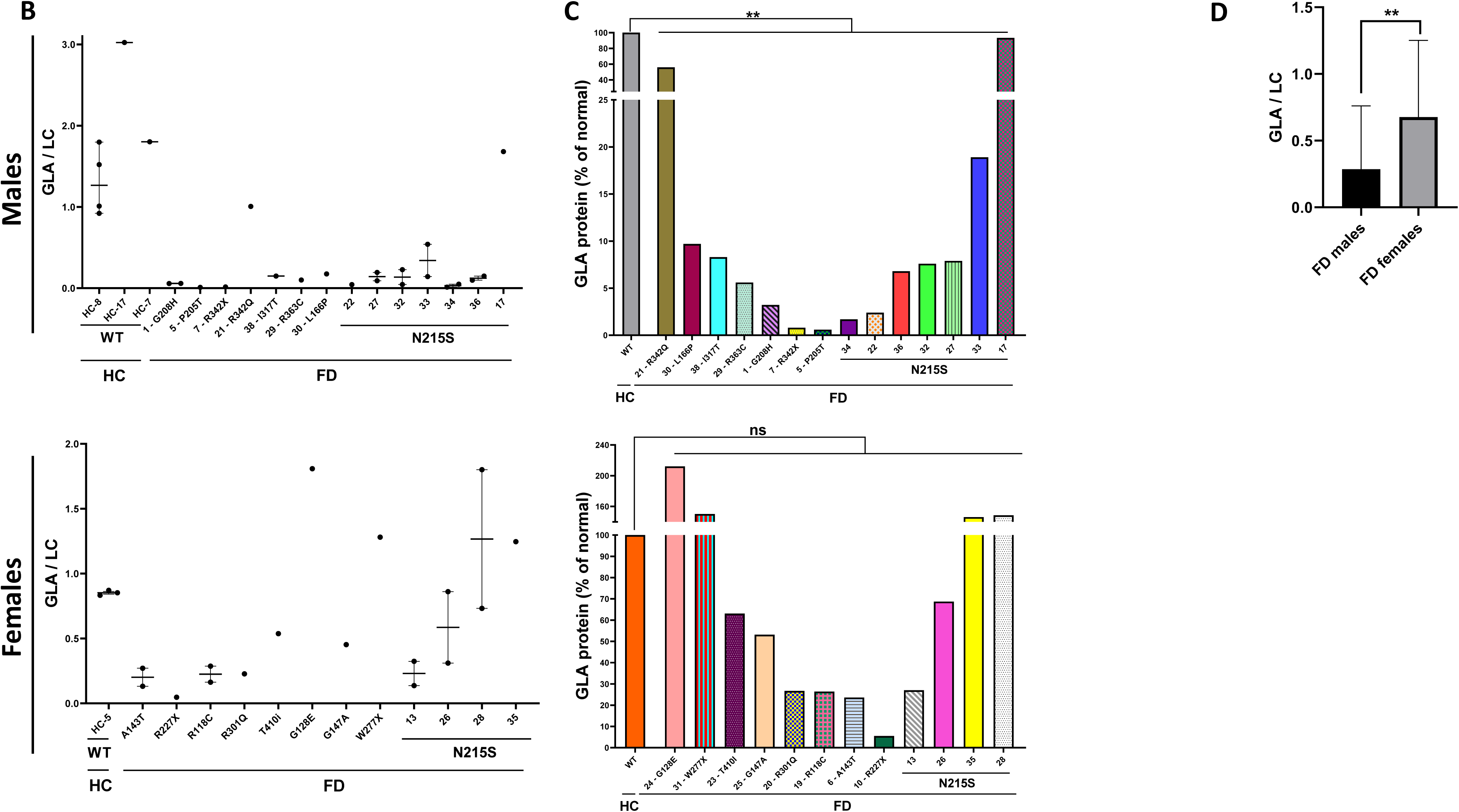
Quantification of α-galactosidase A in mononuclear cells from control and participants with Fabry disease by western blotting analysis. **(A)** Cell lysates containing 10 ug of protein prepared from peripheral blood mononuclear cells (PBMCs) of patients with FD and from healthy controls (HC) were subject to SDS-PAGE and interaction with anti - α-galactosidase A (GLA; 1.3 ng/ul, Bio-techne) antibody and anti-sodium potassium ATPase antibody (0.1 ng/ul, Abcam) antibody, as a loading control (LC). **(B)** For total GLA protein quantification, GLA band intensity in each lane was divided by that of LC in the same lane. **(C)** The median obtained for HCs was set as 100%. Data presented in median and interquartile range; n= 1 to 2 per FD patient, HC= 3 males (n=1, n=1, n=4) and 1 female (n=3). **= p< 0.01, ns= not significant.

All females and most male participants with FD showed two bands corresponding to both forms of GLA protein but with varying intensities, as previously reported (Lemansky et al., 1987). The exception were male patients number 7, 1, and 29, carrying the mutations R342X and G208H, R363C, respectively (fig. 4A). Even in the longest exposure times, only one band was detected for these patients (Supplementary fig. 1). For participant number 22 only 1 band at 50 kDa was detected in this assay, but this was only repeated once. While for most male patients both forms were detectable in the same blot (like in the HCs), for participants 5 and 17 the mature 46 kDa form was only detectable in longer exposure times, hampering its quantification (of note, n= 1 for each; Fig. 4A).

A significant positive correlation was found between total GLA protein levels and GLA activity (males: r2= 0.78 p< 0.0001, females: r2= 0.76 p= 0.008, fig. 5 and supplementary fig. 4). However, as data above suggests the 50 kDa precursor could be driving the enzyme activity measured in the in vitro GLA activity assay (fig. 2), this was assessed for participants with FD too. To that end, both bands were quantified relative to the loading control on the same lane and plotted against their corresponding GLA activity (fig. 6B). A significant correlation was found for female patients 50kDa precursor (r2= 0.69 p= 0.02, fig 6B), indicating that higher amounts of this form corresponded with higher levels of GLA activity, like control participants. Males showed a similar trend (r2= 0.41 p= 0.16, fig 6B) but had significantly less amount of 50 kDa precursor than females (0.12 vs 0.45, p= 0.04, fig. 6A). For both sex groups, amounts of the 46 kDa form were similar and correlation analyses demonstrated negative trends between the mature form and total GLA activity (females: r2= -0.53 p=0.10 and males: r2= - 0.48 p=0.24, fig 6B). No differences between precursor or mature form amounts were found between non-N215S and N215S males.

**Fig. 5.**
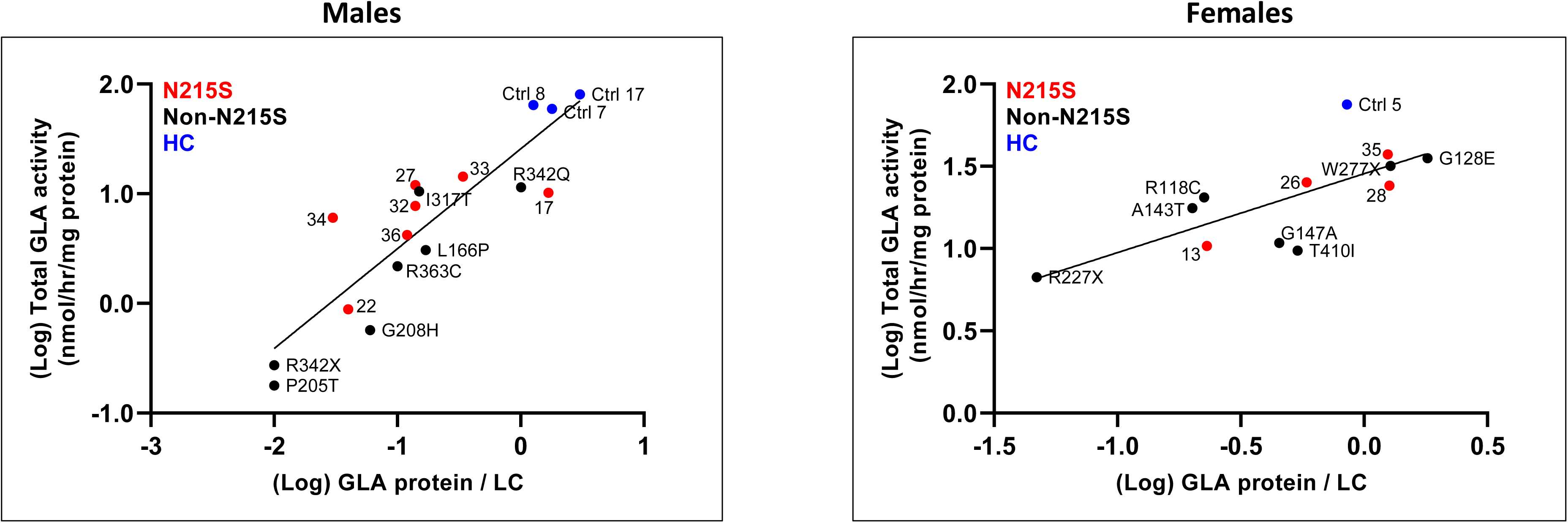
Relation between total α-galactosidase A protein by western blot and α-galactosidase A activity.

**Fig. 6.**
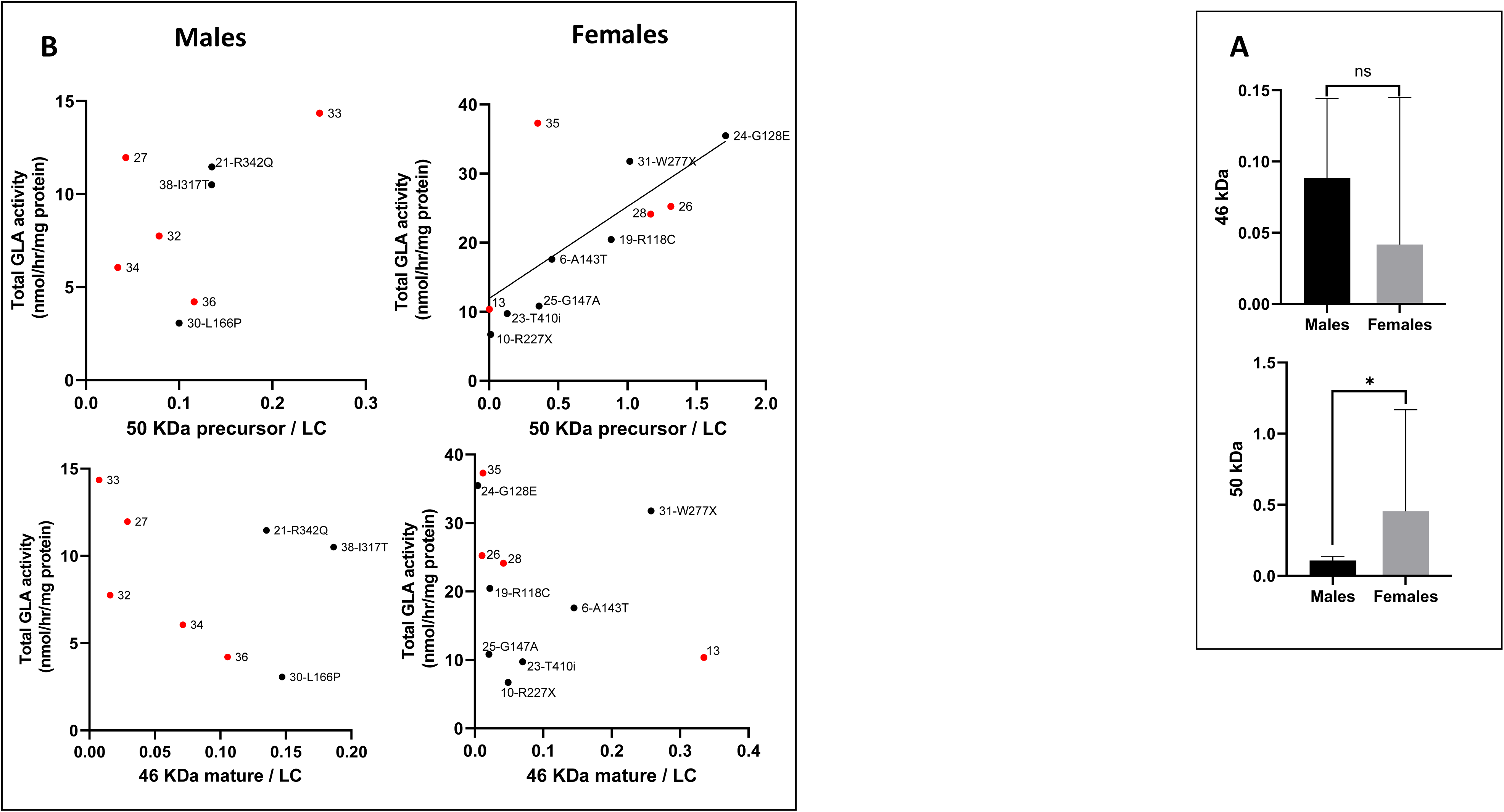
(A) Total amounts of α-galactosidase A (GLA) forms per group. (B) Correlation analysis between GLA forms and residual enzyme activity. To determine the enzymatic activity of the 50 kDa precursor each band was quantified relative to the loading control (LC) in the same lane and plotted against the GLA activity obtained for that subject. Males n= 8 (participants number 1, 5, 7, 17, 22, and 29 were not included). Females n= 11 (participants number 20 was not included as no enzyme activity data was available). Healthy controls (HC) n= 4, 3 males and 1 female.

In terms of clinical severity, for males neither GLA protein or activity showed a correlation with severity scores, GFR, or LVMI. The only significant correlations found through these analyses were between plasma lyso-Gb3 and AASS (p= 0.001, n= 16, fig. 7A and supplementary fig. 5C), and between age and the MSSI and with GFR (MSSI: p= 0.0007, n= 19; GFR: p= 0.0005, n= 18; fig. 7A). Females did show a significant association between GLA activity, the MSSI and GFR (MSSI: p= 0.02, n= 12; GFR: p= 0.01, n= 12; fig. 7A and supplementary fig. 5B), and both overall severity scores correlated with levels of plasma lyso-Gb3 (MSSI: p= 0.04, n= 8; AASS: p= 0.005, n= 8, fig. 7A and supplementary fig. 5B). While females’ GLA activity and plasma lyso-Gb3 correlation was not significant, there was a trend between these (p= 0.07, n= 5; fig.7A and supplementary fig. 5B). This group also showed a strong significant correlation between GLA protein and kidney function and with age (GFR: p= 0.01, n= 12; age: p= 0.02, n= 12; fig. 7A and supplementary fig. 5B). GFR and age were also significantly associated (p< 0.0001, n= 17; fig. 7A).

**Fig. 7A.**
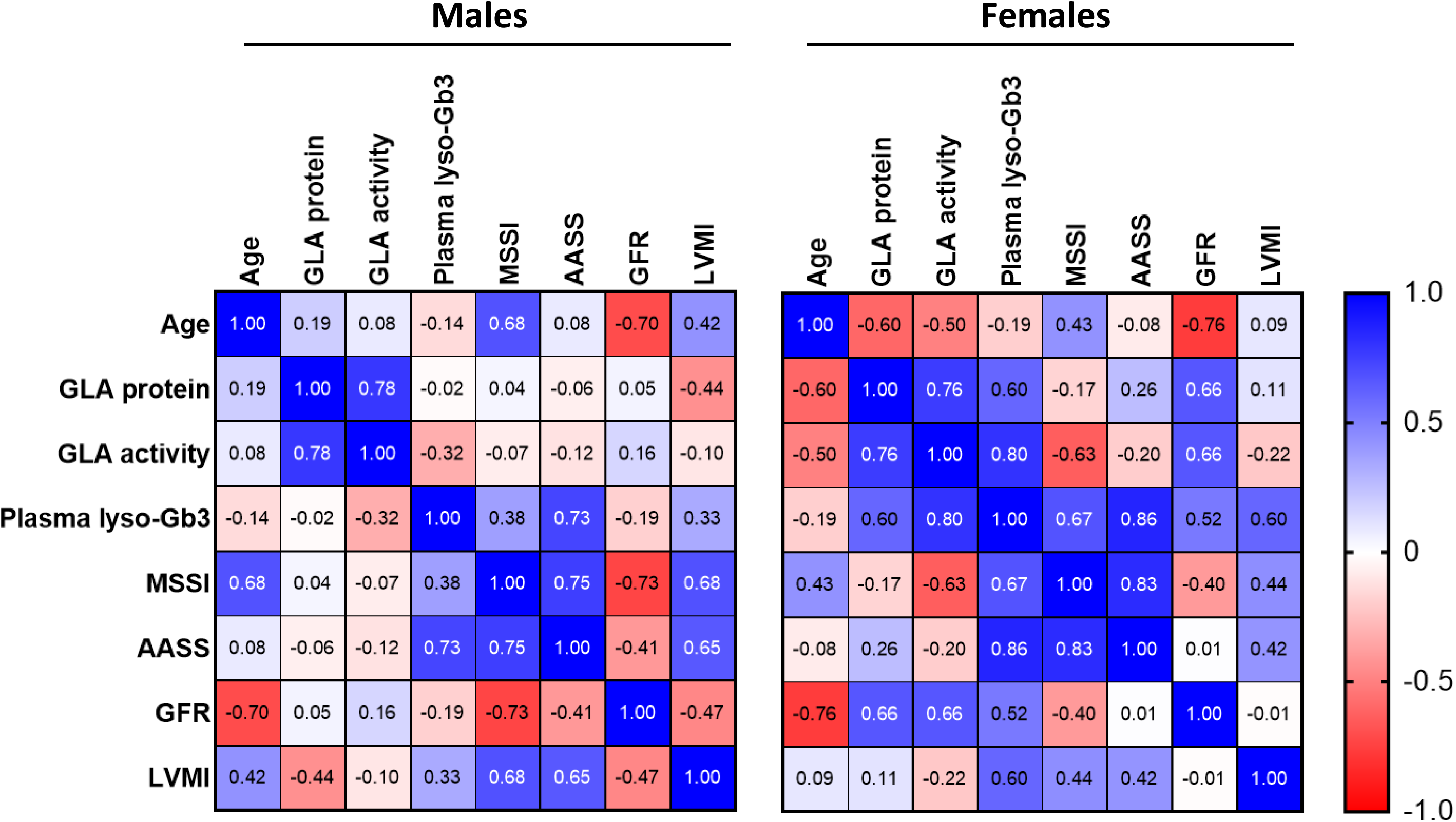
Spearman’s rank correlation matrices between clinical outcomes, GLA activity, protein, age, and plasma lyso-Gb3. The colours represent correlation coefficients, indicating the strength and magnitude of the correlation.

After stratifying male individuals according to their GLA genotype, opposing trends between GLA protein and clinical outcomes became apparent. Interestingly, both groups displayed indirect relations between GLA protein and LVMI, suggesting lower LVMI magnitudes correspond with higher levels of GLA protein. Additionally, the non-N215S group demonstrated associations between GLA activity and MSSI, AASS, LVMI (fig. 7B). The only statistically significant correlations were between age and clinical severity for both genotype groups (N215S: MSSI: p< 0.0001, n= 9; AASS: p= 0.04, n= 9; GFR: p= 0.007, n= 9; LVMI: p= 0.02, n= 6; non-N215S: MSSI: p< 0.008, n= 10; GFR: p= 0.005, n= 9; fig. 7B).

**Fig. 7B.**
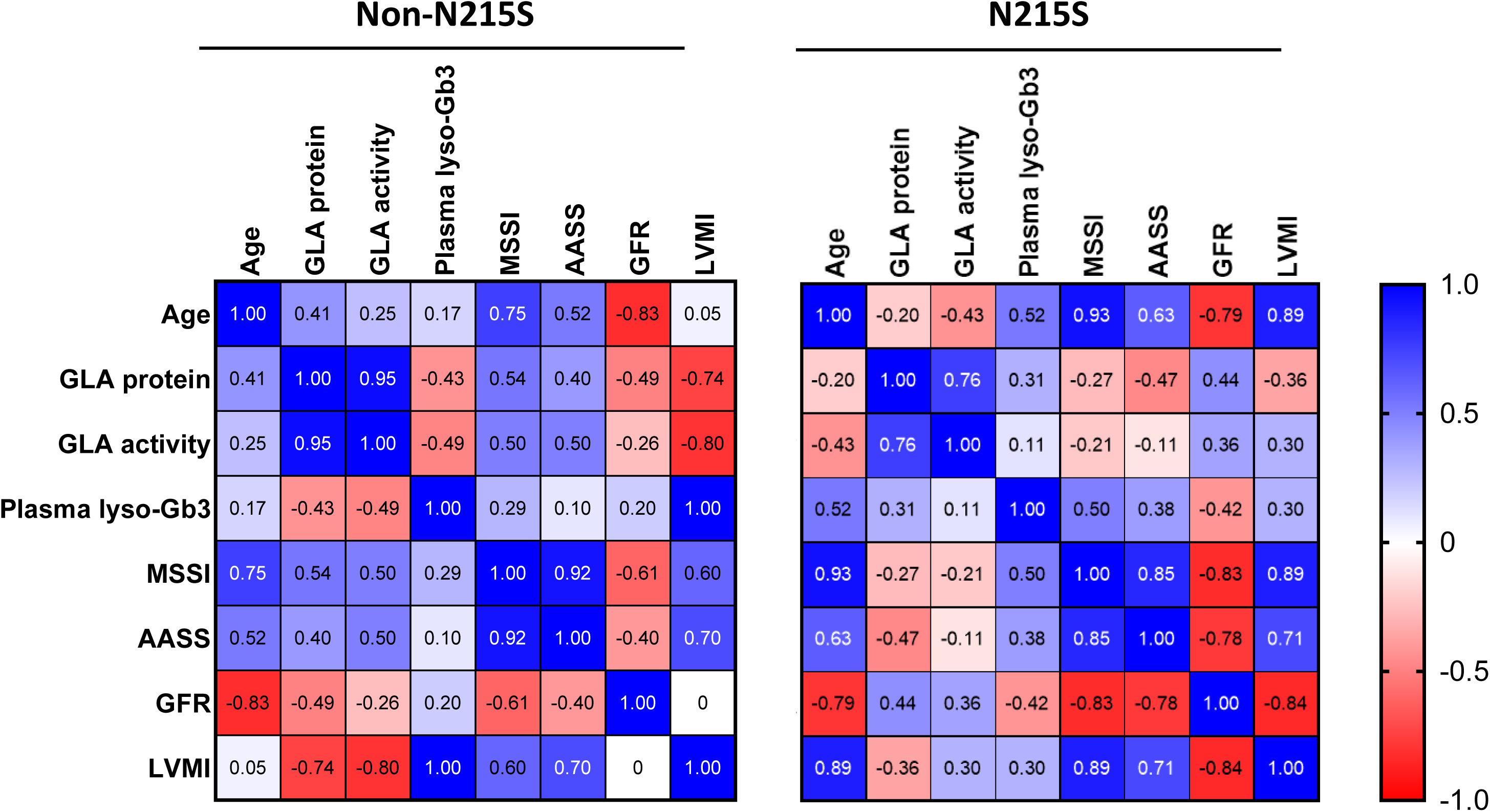
Spearman’s rank correlation matrices between clinical outcomes, GLA activity, protein, age, and plasma lyso-Gb3 for males N215S and non-N215S. The colours represent correlation coefficients, indicating the strength and magnitude of the correlation.

#### 3.3.3 GLA protein forms in fibroblasts and PBMCs from controls and subjects with FD: endo-H assay

GLA protein is synthesised as a 50 kDa precursor which is processed in the lysosomal lumen into the mature lysosomal form of 46 kDa by partial proteolysis, and harbours three N-glycosylation sites: 139, 192 and 215. While the latter two are mostly composed of oligo mannose carbohydrate, the N139 site only contains complex saccharides (Gal et al., 2011; Lee et al., 2003; Matsuura, Ohta, Ioannou, & Desnick, 1998). During processing, these carbohydrates interact with ER resident lectin chaperones, and their composition serves as a folding tag, indicating the folding status of the polypeptide. If the protein fails to attain its proper folding state, the N-glycans will not undergo processing and will instead retain a high-mannose composition and the protein will remain in the ER (Shenkman & Lederkremer, 2019). Therefore, to understand if the variation in GLA protein forms seen for Fabry patients relates to differences in ER processing, whole cell lysates were treated with the endo-H enzyme, able to cleave high-mannose N-glycans (Maley, Trimble, Tarentino, & Plummer, 1989; Trimble & Tarentino, 1991).

In all lysates, GLA protein showed sensitivity to endo-H cleavage indicated by the appearance of one band of about 41 – 42 kDa (fig 8A and 9A, all positive lanes), suggesting that in all subjects there was an ER retained endo-H sensitive form of the protein. The 46 kDa form of GLA protein was the most intense in those lanes corresponding to HC lysates after endo-H digestion (HC positive lanes, fig 8A and 9A), in agreement with data reported in the literature (Braunstein et al., 2020; Brogden et al., 2017; Hamanaka et al., 2008; Ioannou, Zeidner, Grace, & Desnick, 1998). After quantification, results showed that in HC cells about 100% of GLA protein was endo-H resistant, indicating that most of the protein passed the mid-Golgi. Conversely, a median of 13% (5.2% – 70.7%, p= 0.03, fig 9B) of GLA protein in male Fabry patients was endo-H resistant, suggesting that a significant fraction was not lysosomal. While female subject showed higher levels of the 46 kDa form, males had higher levels of the 50 kDa immature form (fig. 9C). No difference between male patients with the N215S genotype and those with other missense mutation were found in total levels of GLA protein endo-H resistant form. Female patients with missense mutations showed an endo-H resistant fraction of 4.8% to 119.9%, with no difference from that of the gender matched control (fig. 9B). Only two female patients with a nonsense mutation took part in the study, patient 10 harboured the R227X mutation and her endo-H resistant fraction was of 4.8% and patient 31 (W277X), who demonstrated an endo-H resistant fraction of 104.8% of HC. While the former showed a total GLA protein of 5.5% and a GLA activity of 11.6% of normal, the latter had a total GLA protein of 150.3% with a REA of 55% from HC. Of note, patient 10 was 39 years old when the sample was taken and had an AASS of 0.7, whereas patient 31 was 68-year-old and her AASS was of 3.2.

**Figure 8.**
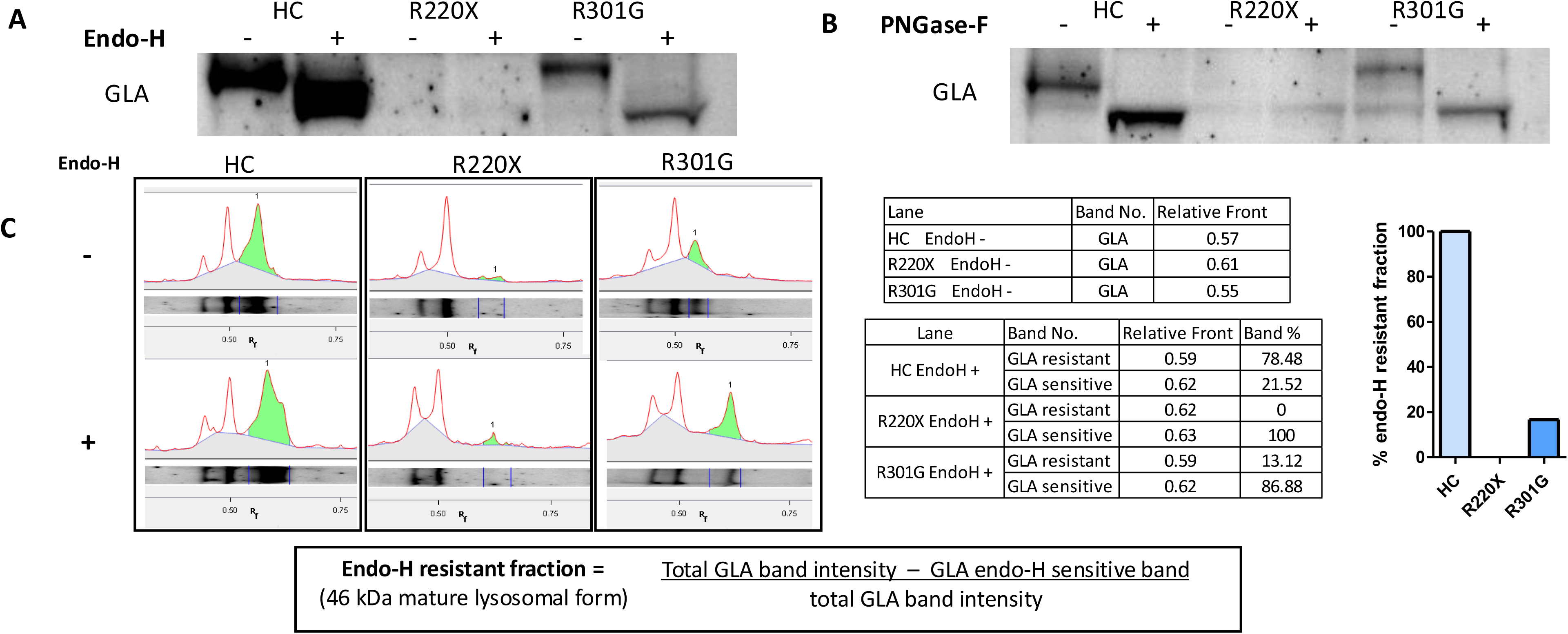
Endoglycosidase H resistance of α-galactosidase A in control and FD fibroblasts. Cell lysates with the same amount of protein were prepared from fibroblast derived from 2 subjects with GLA variants: Arg220Ter (R220X) and Arg301Gly (R301G) and an apparently healthy individual (HC) GLA variant. Samples were subjected to **(A)** endoglycosidase H (endo-H) and **(B)** Peptide:N-glycosidase F (PNGasa-F), digestion. In parallel, undigested control samples were prepared by using water instead of the enzymes (endo-H – lane). After overnight incubation at 37°C, samples were subject to SDS-PAGE and interaction with anti - α-galactosidase A (GLA; 1.3 ng/ul, Bio-techne) antibody. **(C)** Densitometric analysis and quantification of endo-H resistant fraction: blots were scanned, and the intensity of each band was measured. GLA resistant fraction was calculated by dividing the difference between the intensity of the total GLA band (in the endo-H + lane) and the intensity of the endo-H sensitive fraction (in the endo-H + lane) by the intensity of the total GLA band in the same lane. The value obtained for the HC was set as 100%. N= 1 for each group.

**Fig. 9.**
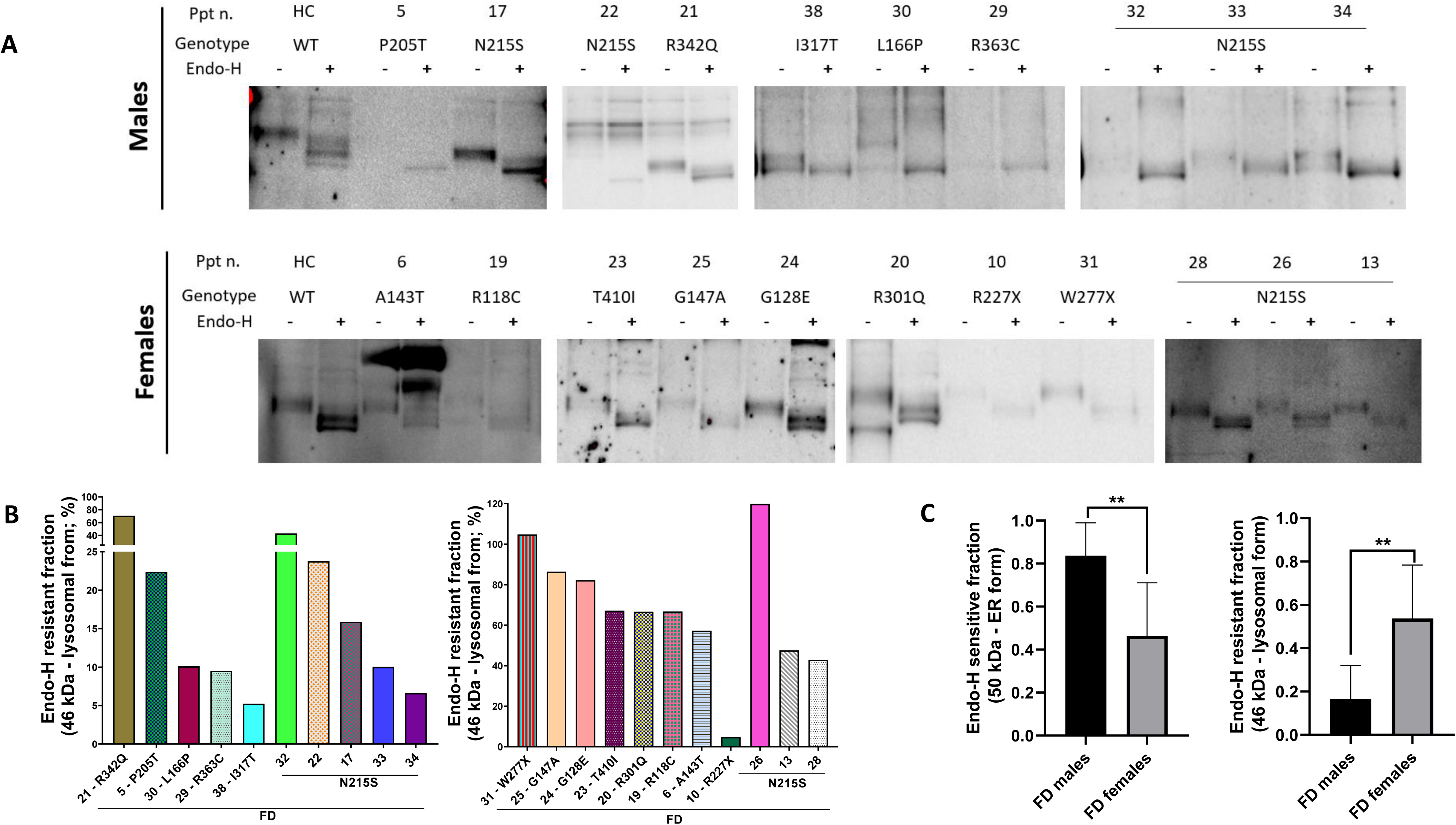
Endo-H resistance of α-galactosidase A in mononuclear cells from control and participants with Fabry disease. **(A)** Cell lysates containing the same amount of protein prepared from peripheral blood mononuclear cells (PBMCs) of patients with Fabry disease (FD) and from healthy controls (HC) were subjected to endo-H digestion and western blot analysis. In parallel, undigested control samples were prepared by using water instead of the enzymes (endo-H – lane). After overnight incubation at 37°C, samples were subject to SDS-PAGE and interaction with anti - α-galactosidase A (GLA; 1.3 ng/ul, Bio-techne) antibody. **(B)** To determine the endo-H resistant fraction, the blots were scanned, and the intensity of each band was measured. GLA resistant fraction was calculated by dividing the difference between the intensity of the total GLA band (in the endo-H + lane) and the intensity of the endo-H sensitive fraction (in the endo-H + lane) by the intensity of the entire amount of the GLA in the same lane. The median obtained for HCs was set as 100%. Mann-Whitney tests between HC and FD patients and **(C)** between male and female patients (n=1 per participant) were done to study differences in endo-H resistant and sensitive fractions.

To shed light on the GLA protein form responsible for the enzyme activity measured in the enzyme activity assay employed in this study, each form (ER and lysosomal) were analysed against the GLA activity obtained for the corresponding participant (fig. 10). Females displayed negative association between total GLA activity and the endo-H sensitive form, suggesting that higher levels of this form corresponded with lower GLA activity levels (r2= -0.56, p= 0.05, fig. 10). In contrast, their endo-H resistant fraction exhibited a positive association with total GLA activity (r2= 0.59, p= 0.04, fig. 10). While data for males exhibited similar trends, these associations were not statistically significant (fig. 10).

**Fig. 10.**
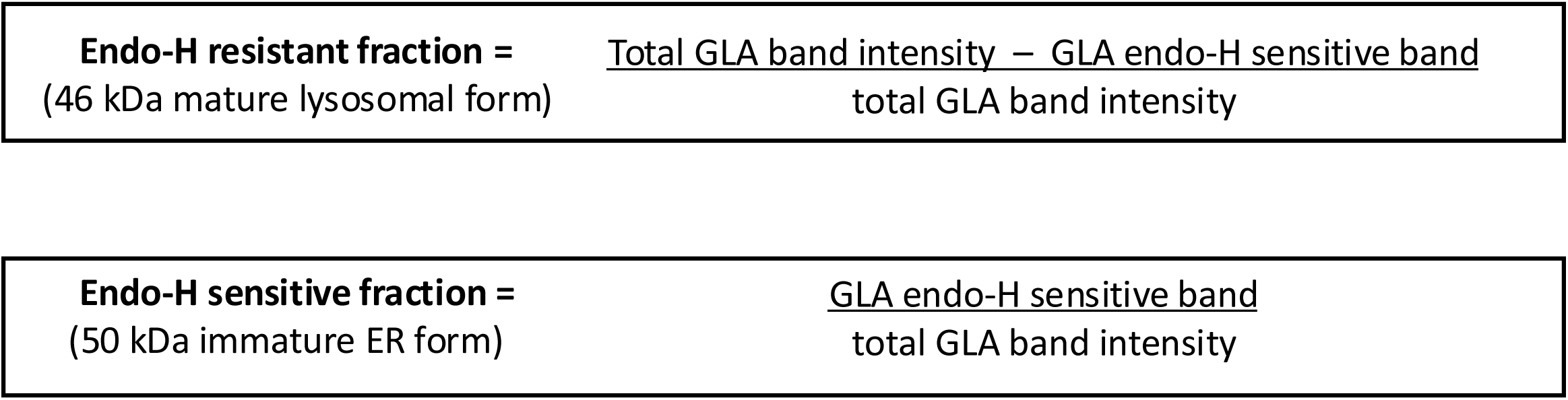
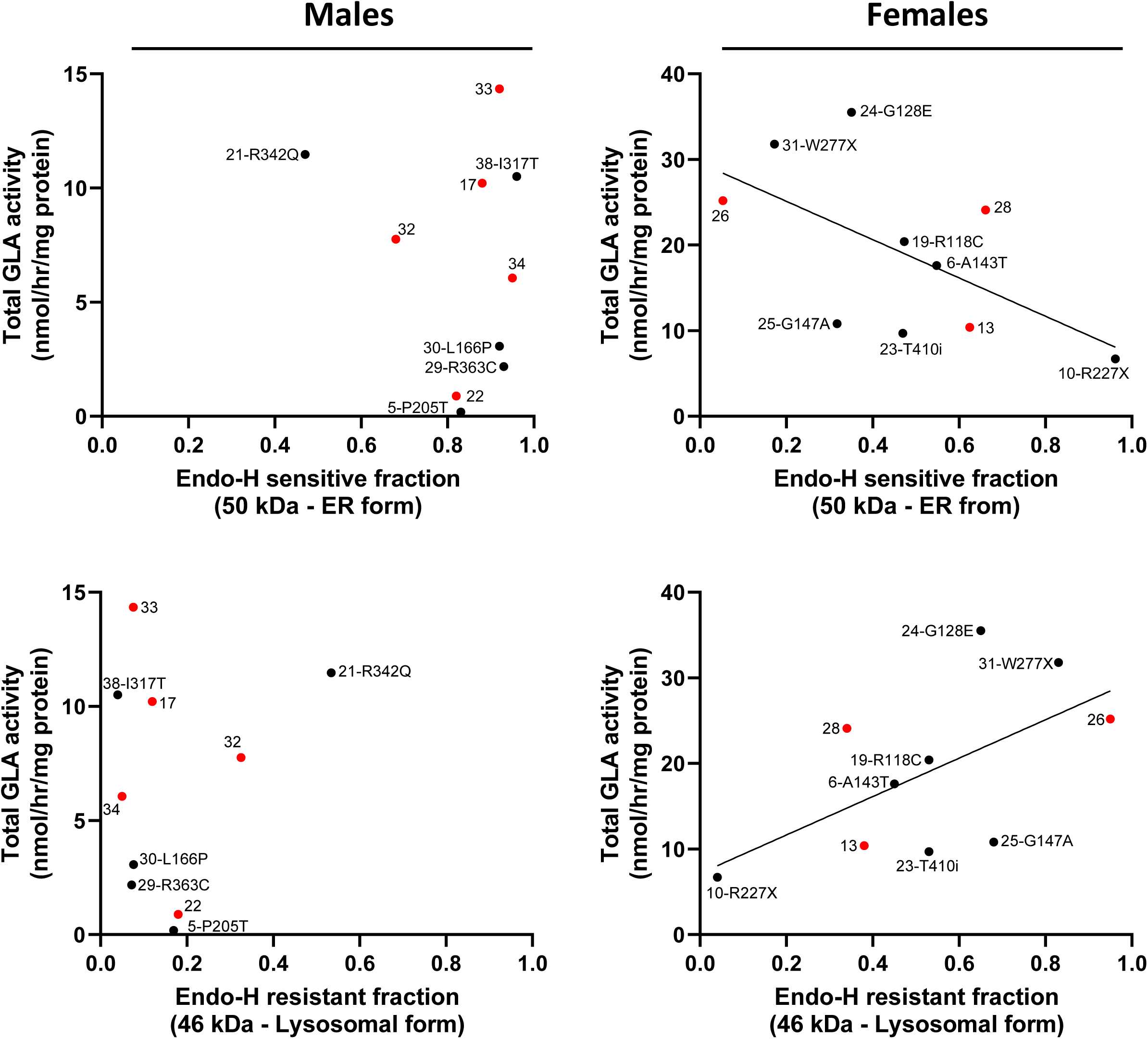
Correlation analysis between α-galactosidase A endoglycosidase H forms and residual enzyme activity. To determine the enzymatic activity of the α-galactosidase A (GLA) endoglycosidase H (Endo-H) sensitive and resistant forms, the intensity of the endo-H sensitive band was measured (in the endo-H + lane) and divided by the intensity of the entire amount of the GLA in the same lane. Then each form was plotted against the enzyme activity obtained for that patient. Males n= 10 and females n= 10 (participants number 20 was not included as no enzyme activity data was available).

In terms of clinical severity, there were similar trends between the endo-H forms and LVMI, but these were only statistically significant for females. While higher levels of the sensitive ER form corresponded with lower values of LVMI, the opposite was found for the resistant lysosomal fraction, which showed a positive association with LVMI (fig. 11 and 12). Conversely, opposing trends between endo-H forms and age became apparent among sex groups. Higher levels of the resistant lysosomal GLA form were found in older males, whereas higher levels of the sensitive ER GLA form were found in the younger group (fig. 11 and 13). To assess the available GLA activity within the lysosomal lumen, the 46 kDa lysosomal form fraction was multiplied by the GLA activity obtained for the same participant. This parameter appeared to relate to clinical and biochemical outcomes only in females (fig. 11 and 14).

**Fig. 11.**
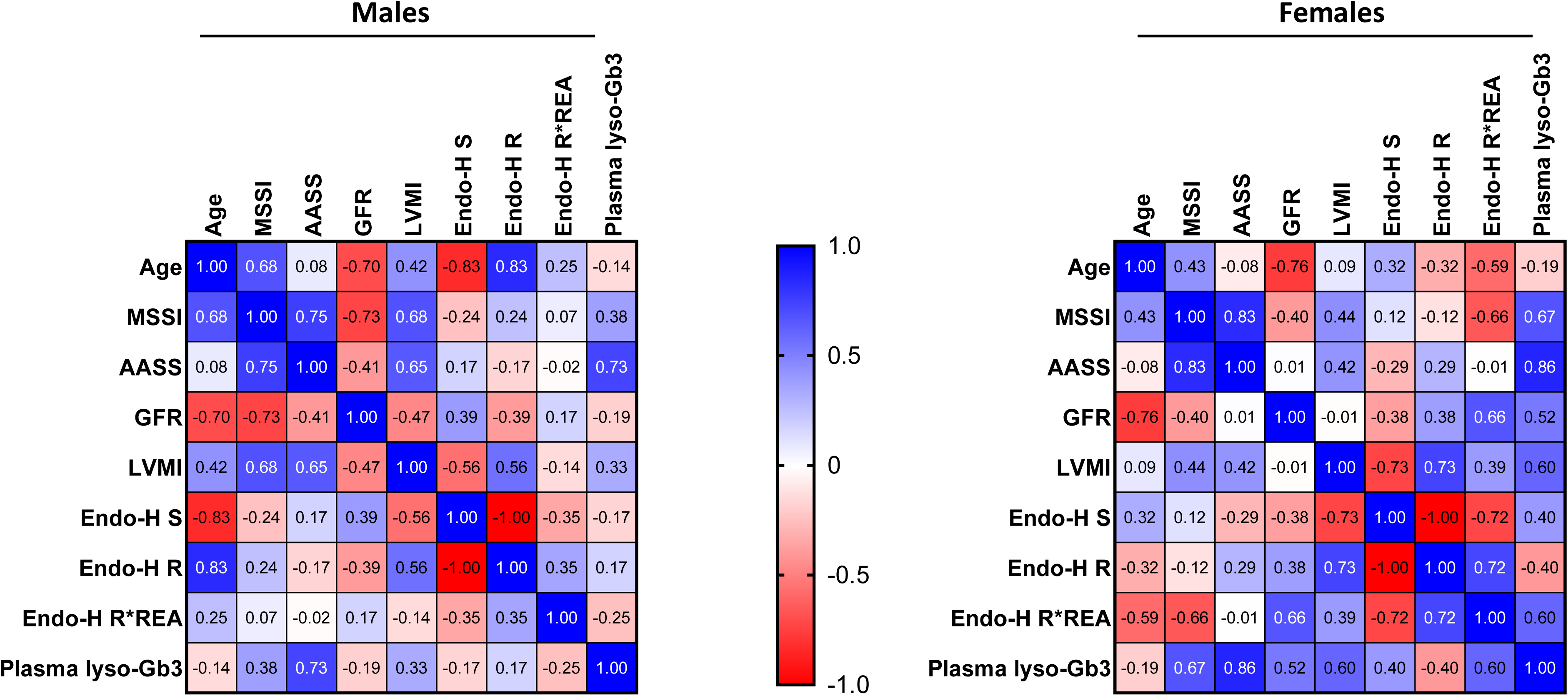
Spearman’s rank correlation matrices between clinical outcomes and GLA protein forms, lysosomal GLA activity, and plasma lyso-Gb3. Lysosomal GLA activity was defined as the total GLA activity times the endo-H resistant (endo-H R) coefficient for the same participant (endo-H R*REA). Endo-H sensitive (Endo-H S). The colours represent correlation coefficients, indicating the strength and magnitude of the correlation.

**Fig. 12.**
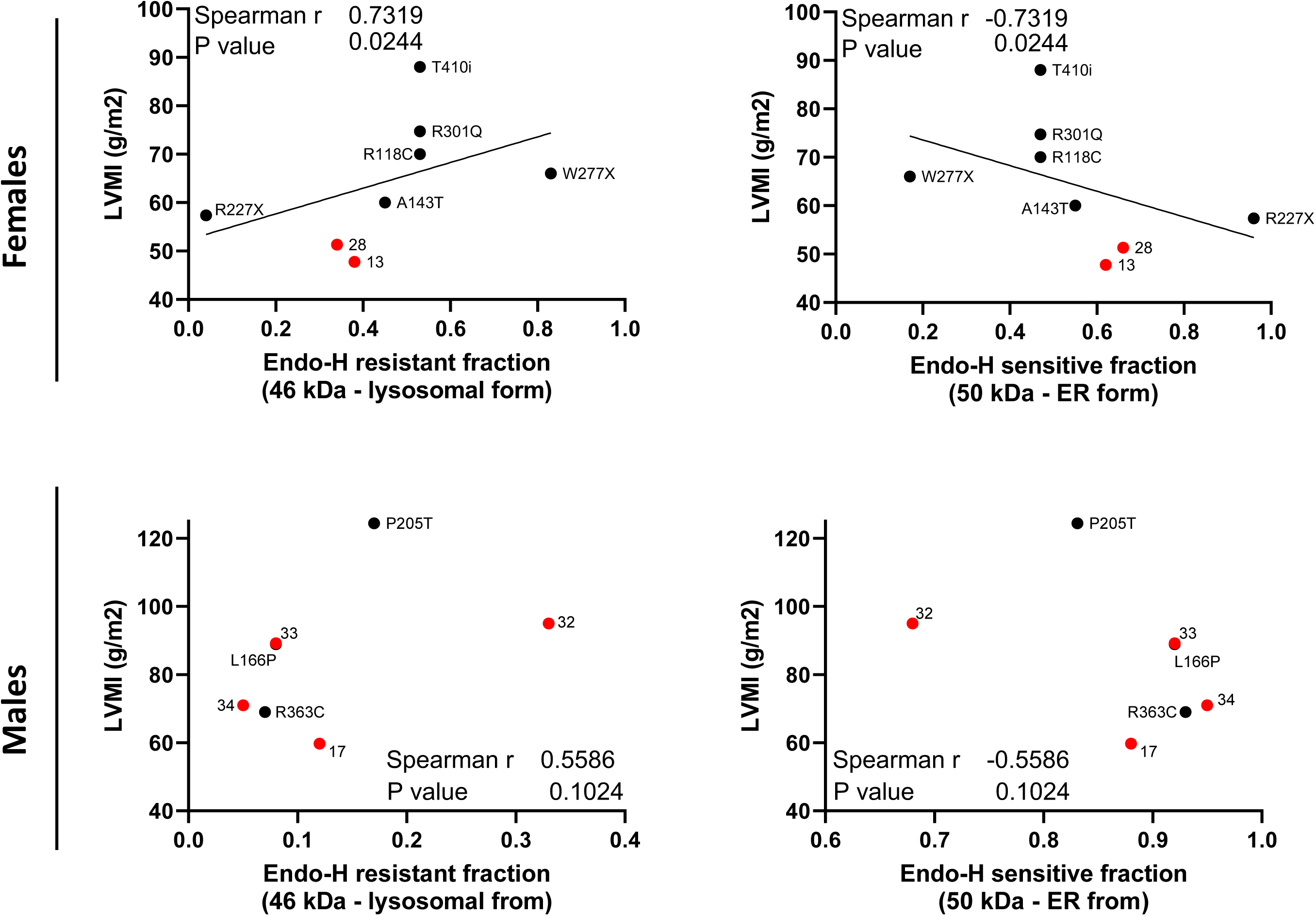
Correlation analyses between GLA endo-H forms and LVMI. In red are those subjects with the N215S genotype.

**Fig. 13.**
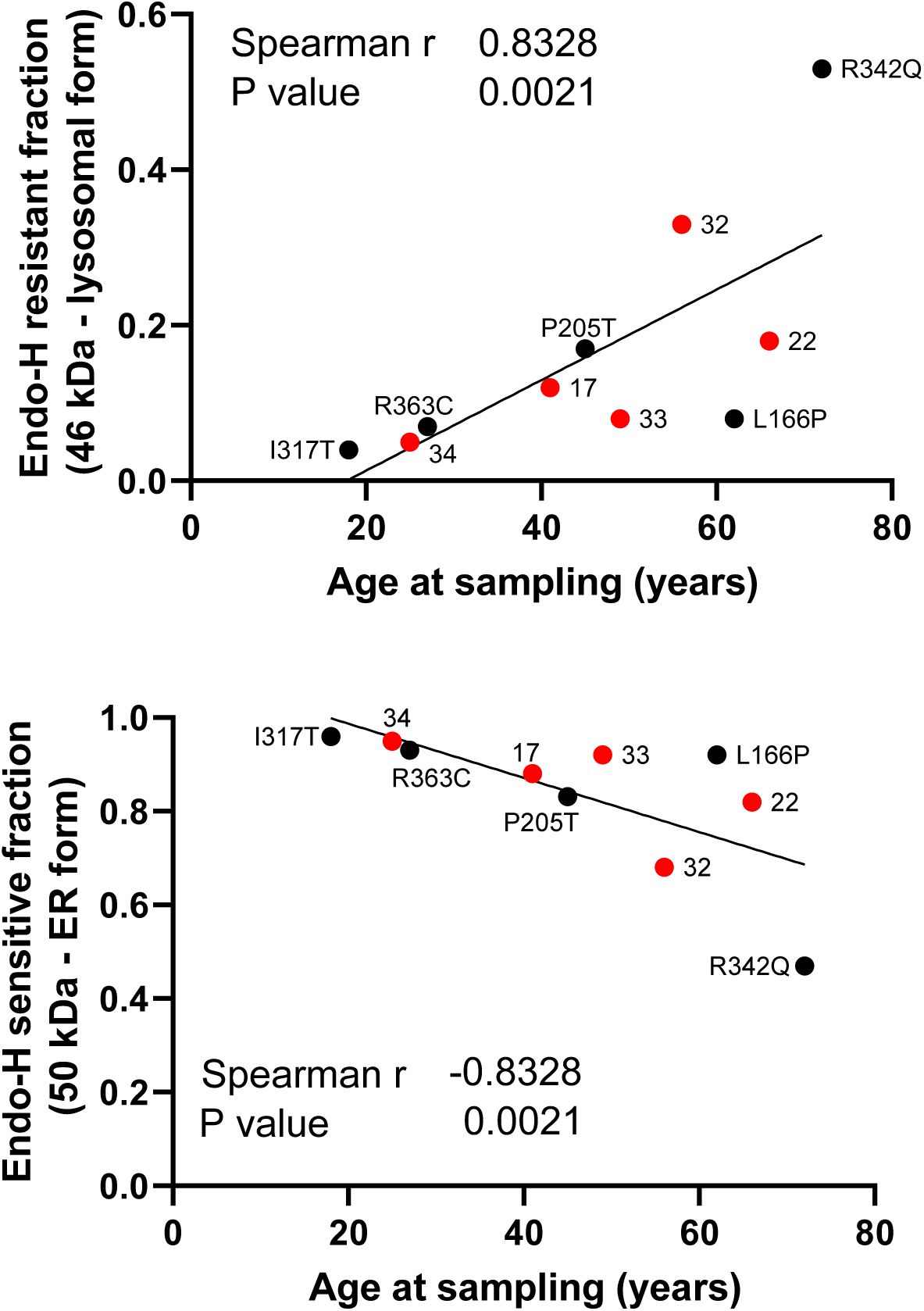
Correlation analyses between GLA endo-H forms and age in males.

**Fig. 14.**
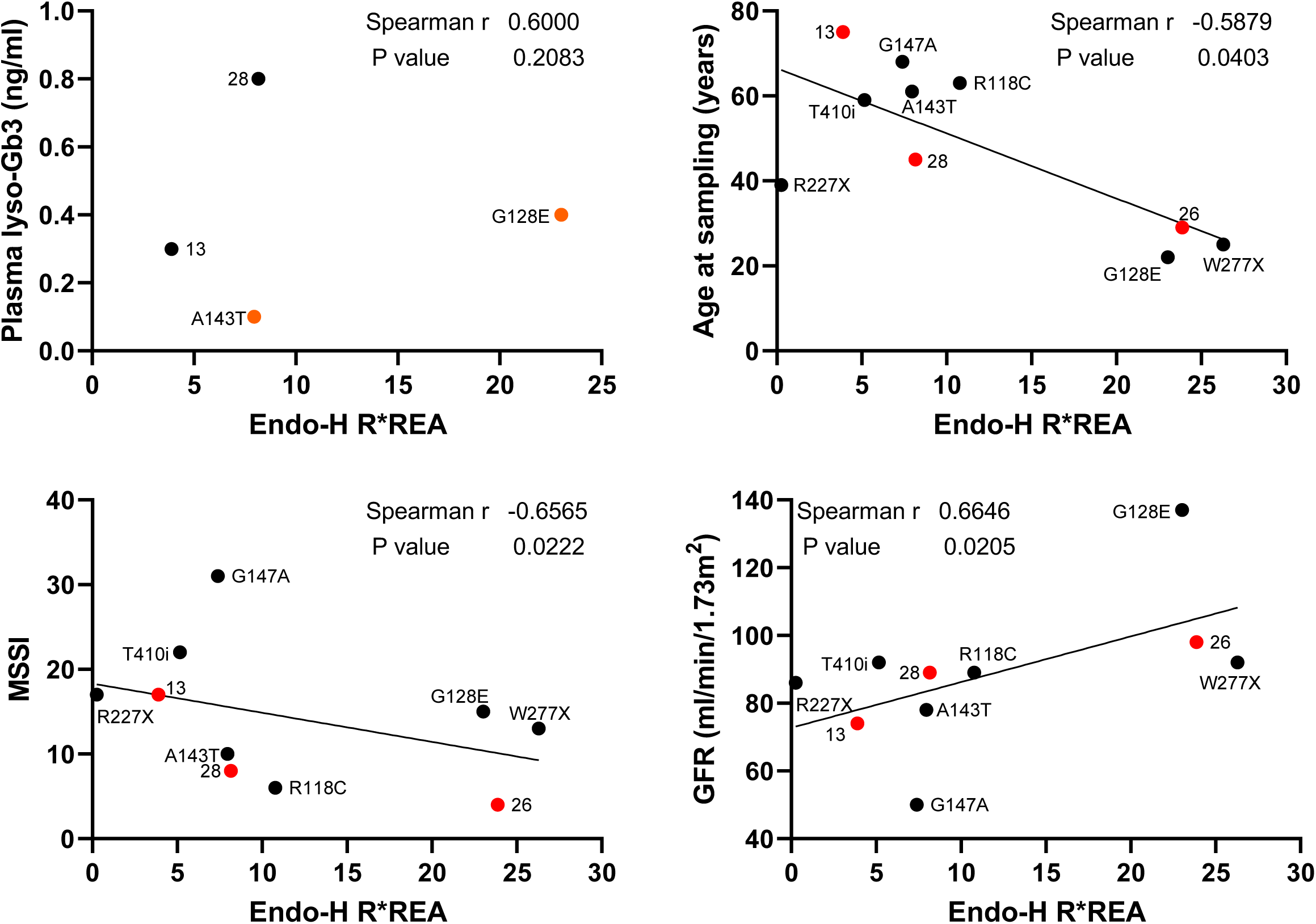
Correlation analyses of lysosomal GLA activity with clinical and biochemical outcomes for females. Lysosomal GLA activity was defined as the total GLA activity times the endo-H resistant coefficient for the same participant (endo-H R*REA).

#### 3.3.4 Plasma lyso-Gb3 data

Lyso-Gb3 data was available for 16 males and 8 females, of which 8 and 5 participants were on treatment at the time the sample was taken, respectively. As previously reported, males had higher levels than females (6.4 vs 1.0 ng/ml, p= 0.004, supplementary fig. 5A), but lyso-Gb3 plasma levels only showed a trend with GLA activity and protein in females (fig. 7A and supplementary fig. 5B). In terms of clinical manifestations, females showed all positive associations between plasma lyso-Gb3 levels and all clinical outcomes i.e., MSSI, AASS, GFR, and LVMI, but only the correlations with the severity scores achieved statistical significance (MSSI: r2= 0.67 p= 0.038, AASS: r2= 0.86 p= 0.005, fig. 7A and supplementary fig. 5B). For males, plasma lyso-Gb3 only showed a correlation with the AASS, also statistically significant (r2= 0.73 p= 0.001, fig. 7A and supporting fig. 5C).

## 4.0 Discussion

To study the relation between GLA activity and GLA protein in FD, fibroblast cell lines and PBMCs from individuals with FD and HC were examined. As most participants were on treatment at the time sample was taken, cells were cultured for 6 days as the compounds half-life are of 3 to 5 hours for PCT (Information, 2023b) and up to 80 minutes for agalsidase alfa (Pastores, 2007) and algasidase beta (considering a 1 mg/kg dose with a mean infusion length of 115 minutes)(Information, 2023a). After this wash out period, HC’s PBMCs showed higher levels of GLA activity and protein, however, the former seemed to increase at the expenses of 50kDa form of the protein. This experiment also suggested that this 50 kDa precursor form was driving the activity measured in the invitro enzyme assay employed. When looking at male patients’ data, no correlation was observed between GLA activity and endo-H resistant (46 kDa lysosomal) fraction. Although this was not the objective of their analysis, Riillo et al similarly reported WT GLA activity levels but significantly higher ER retention for the S126G variant in an in vitro model of the disease (Riillo et al., 2023). Given that they also employed whole cell lysates and the same assay to measure GLA activity, together these finding could question whether these results obtained through this in vitro assay accurately reflect the available GLA activity inside patients’ lysosomes.

Significant correlations were found between GLA activity and endo-H fractions in females, suggesting higher activity levels corresponded to a higher endo-H resistant fraction (46 kDa lysosomal) but lower levels of endo-H sensitive fraction (50 kDa ER form). Additionally, female patients exhibited higher GLA activity and protein levels than their male counterparts, and showed significant correlations between clinical severity (MSSI and GFR) and GLA activity, in line with findings from other groups (Arends et al., 2017; Branton et al., 2002; Lukas et al., 2013; Oliveira & Ferreira, 2019). As females also had higher levels of the 46 kDa lysosomal form, these observations could be explained by the impact of the expression of the WT GLA allele due to favourable skewing during Lyonization. This aligns with the concept that skewed X-inactivation is a major factor influencing the severity of clinical expression in Fabry heterozygotes (Beck & Cox, 2019). Conversely, males had higher levels of the 50 kDa ER form, which seems to increase levels in the cellular culture conditions employed in this study. As this form appeared to be enzymatically active in the in vitro assay used, the lack of correlation observed between GLA activity and clinical severity in males could be the consequence of artificial GLA activity results, measured from whole cell lysates, outside of physiological conditions.

Endo-H experiments suggested subjects with FD had relatively higher levels of the immature 50 kDa form of GLA protein (endo-H sensitive ER form), in line with findings of other research teams who reported that most GLA missense mutation result in protein mutants that are retained in the ER (Lenders, Stappers, & Brand, 2020; Yam, Zuber, & Roth, 2005). Interestingly, these forms were associated with LVMI, with higher levels of the 50 kDa form corresponding with lower LVMI magnitudes in females. While for males these endo-H forms did not show associations with clinical outcomes, they showed strong correlations with age, suggesting that higher levels of the sensitive 50 kDa form are found on younger patients, while the mature lysosomal 46 kDa form are found in older ones. N-glycosylation is known to be influenced by aging and can lead to an increase in certain types of glycoforms after the age of 40 years old (Vanhooren et al., 2010). However, this could also reflect a diagnostic bias towards males, who are consequently screened for the disease earlier in life, potentially explaining the higher levels of the immature GLA 50 kDa ER form of the protein in this group only. This is in line not only with findings in FD (Ellaway, 2015), but also with findings in many other conditions, where males seemed to be diagnosed at a younger age (Westergaard, Moseley, Sørup, Baldi, & Brunak, 2019). Another interesting difference between sex groups was observed in the relation between age and LVMI, which was only found for males. However, after stratifying individuals based on their genotype, this association persisted exclusively within the N215S group. Therefore, this observation could be reflecting the fact that individuals with this genotype mostly present with cardiac involvement and have greater overall survival (Lavalle et al., 2018).

Results from this work suggest a link between levels of total GLA protein and severity in Fabry clinical manifestations. For females this link would seem to be (at least partially) via GLA activity, as seen in the significant positive association between levels of endo-H mature 46 kDa lysosomal form and total GLA activity. Additionally, both total GLA activity and lysosomal GLA activity were significantly associated with kidney function and MSSI. For males, no correlation was found between total GLA protein amounts, activity, and clinical outcomes. While the results above could suggest this is due to inadequate enzyme activity assay, unlike in females, lysosomal GLA activity did not exhibit correlation with any of the clinical outcomes assessed.

Differences in total GLA protein between different gene variants have been previously studied by several groups and this work has led to the development of a new therapy for the disease i.e., PCT, which stabilizes the misfolded mutant and facilitates its trafficking to the lysosome (Benjamin et al., 2017). New therapeutic strategies focus on enhancing misfolded protein degradation, for which Seemann et al tested the impact of a variety of drugs that inhibit the proteosome on GLA activity in FD fibroblast. They saw a varying response in terms of GLA activity enhancement among drugs and a significant difference on GLA activity increase between variants regardless of the proteosome inhibitor tested (Seemann et al., 2020).

The findings presented in this study highlight the complex relationship between GLA protein forms, enzyme activity, and clinical outcomes in FD, with distinct differences observed between male and female patients. The observed discrepancies in GLA activity and its correlation with disease severity between sexes underscore the potential influence of factors such as X-inactivation, age, and diagnostic timing. These results enhance our understanding of the molecular mechanisms underlying FD and suggest that current enzyme activity assays may need refinement to more accurately reflect lysosomal function. Further research is needed to explore these differences and to develop therapeutic strategies that consider the nuanced interplay between GLA protein forms and clinical heterogeneity in FD.

## 5.0 Limitations

This is a single-centre cross-sectional study and the laboratory results presented correspond to a single time point sample. Clinical data for all patient was assessed retrospectively and only a subset of patients had lyso-Gb3 levels and LVMI data available for analysis.

Regarding the endo-H assay, this is a specific endoglycosidase that can distinguish between high mannose sugars and a mature N-glycan complex. As proteins are correctly processed and trafficked through the ER and Golgi apparatus, mannoses are removed from the glycans, and the proteins becomes resistant to endo-H digestion. Hence, this reaction differentiates between fully mature glycoproteins and those retained in the ER (Maley et al., 1989). Nevertheless, changes in protein migration on SDS-PAGE could originate from changes in the amino acid sequence of the protein rather than from changes in the glycan. To assess this, cell lysate should also be subjected to Peptide:N-glycosidase F digestion as this enzyme removes all aspargine-linked glycans, allowing to detect the molecular weight of the fully deglycosylated protein (Cuddy & Mazzulli, 2021). Another important control for the endo-H assay is the detection of another endo-H sensitive protein to exclude a general decrease in the number of lysosomal enzymes. For example, β-hexosaminidase A as its mature form has at least one N-glycan complex (Mahuran, 1995; Ron & Horowitz, 2005). In this study, these controls were not systematically assessed in patient samples and thus add an additional layer of confounders.

## 6.0 Conclusion

GLA protein processing appeared to impact clinical manifestations in individuals with FD. While for females this relationship could relate to the expression of the WT allele and therefore relate to substrate levels secondary to residual GLA activity, for males this does not seem to be the case.

## Notes

### Competing Interest Statement

The authors have declared no competing interest.

### Funding

- Royal Free Charity

