## Supplementary figures for "Sex differences in alpha galactosidase protein processing and its impact on disease severity in Fabry disease"

### Slide 1
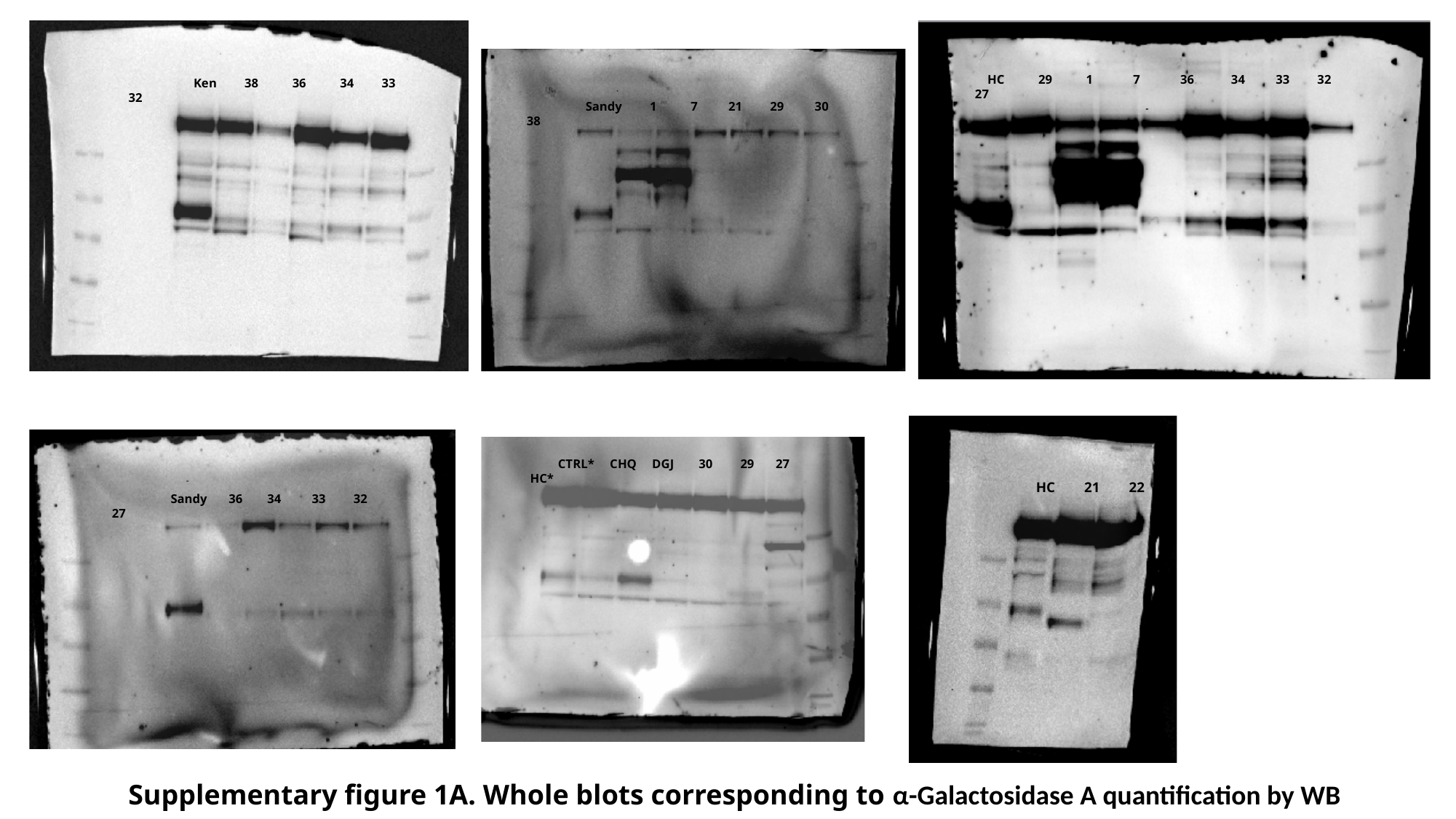

HC 29 1 7 36 34 33 32 27
 Ken 38 36 34 33 32
 Sandy 1 7 21 29 30 38
 CTRL* CHQ DGJ 30 29 27 HC*
 HC 21 22
 Sandy 36 34 33 32 27
Supplementary figure 1A. Whole blots corresponding to α-Galactosidase A quantification by WB

### Slide 2
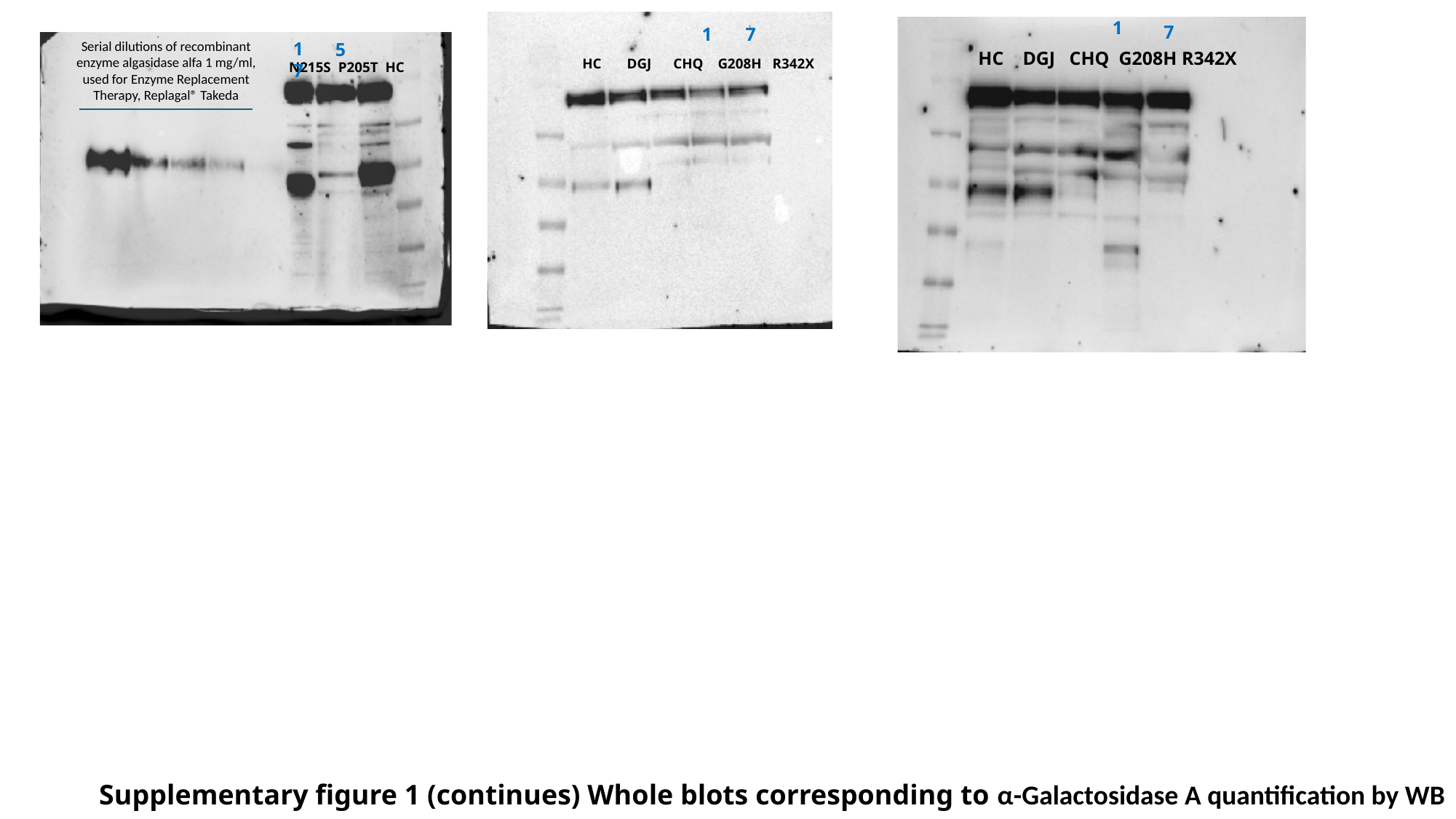

1
7
 HC DGJ CHQ G208H R342X
1
7
 HC DGJ CHQ G208H R342X
17
5
N215S P205T HC
Serial dilutions of recombinant enzyme algasidase alfa 1 mg/ml, used for Enzyme Replacement Therapy, Replagal® Takeda
Supplementary figure 1 (continues) Whole blots corresponding to α-Galactosidase A quantification by WB

### Slide 3
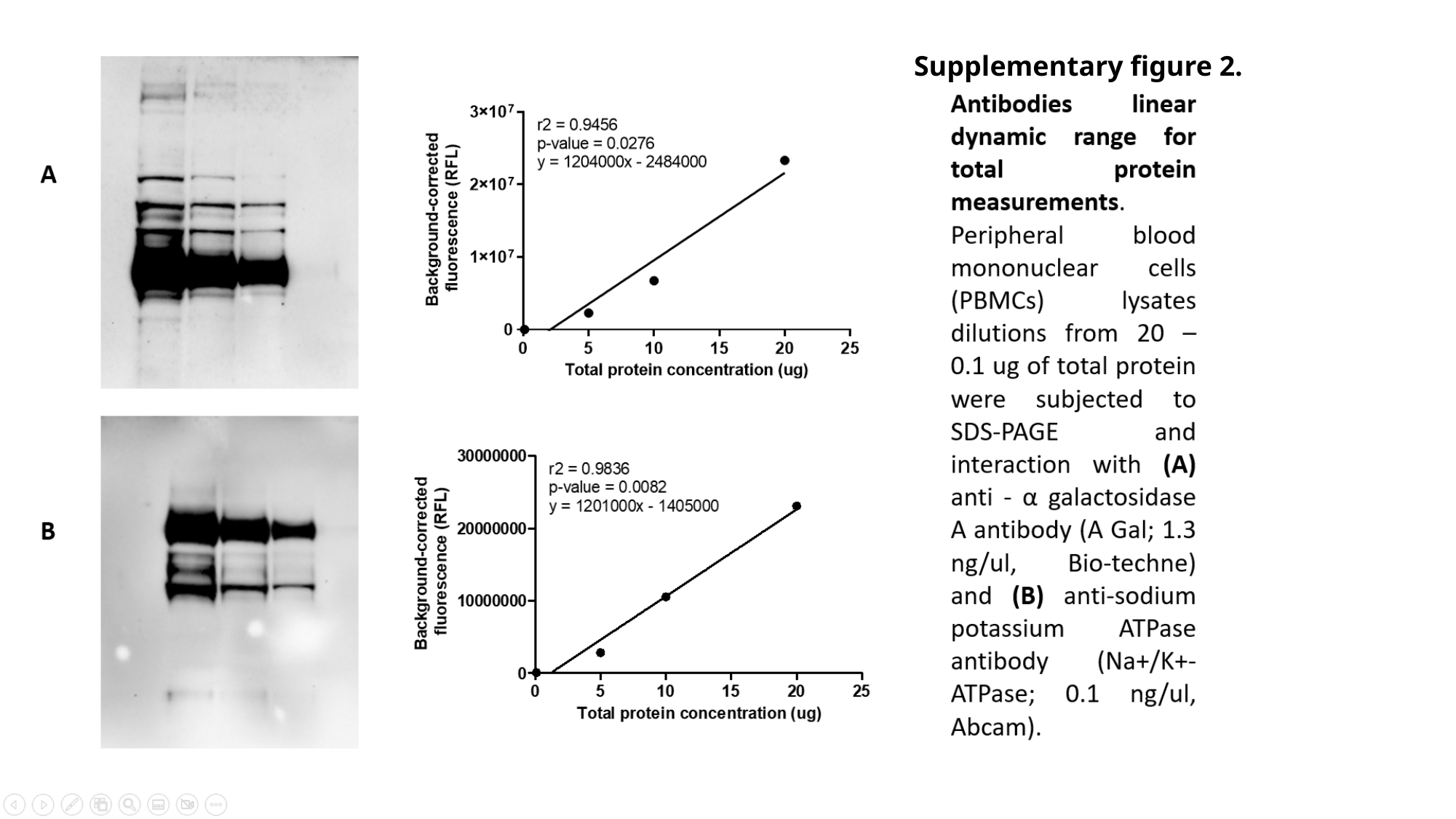

#
Supplementary figure 2.

### Slide 4
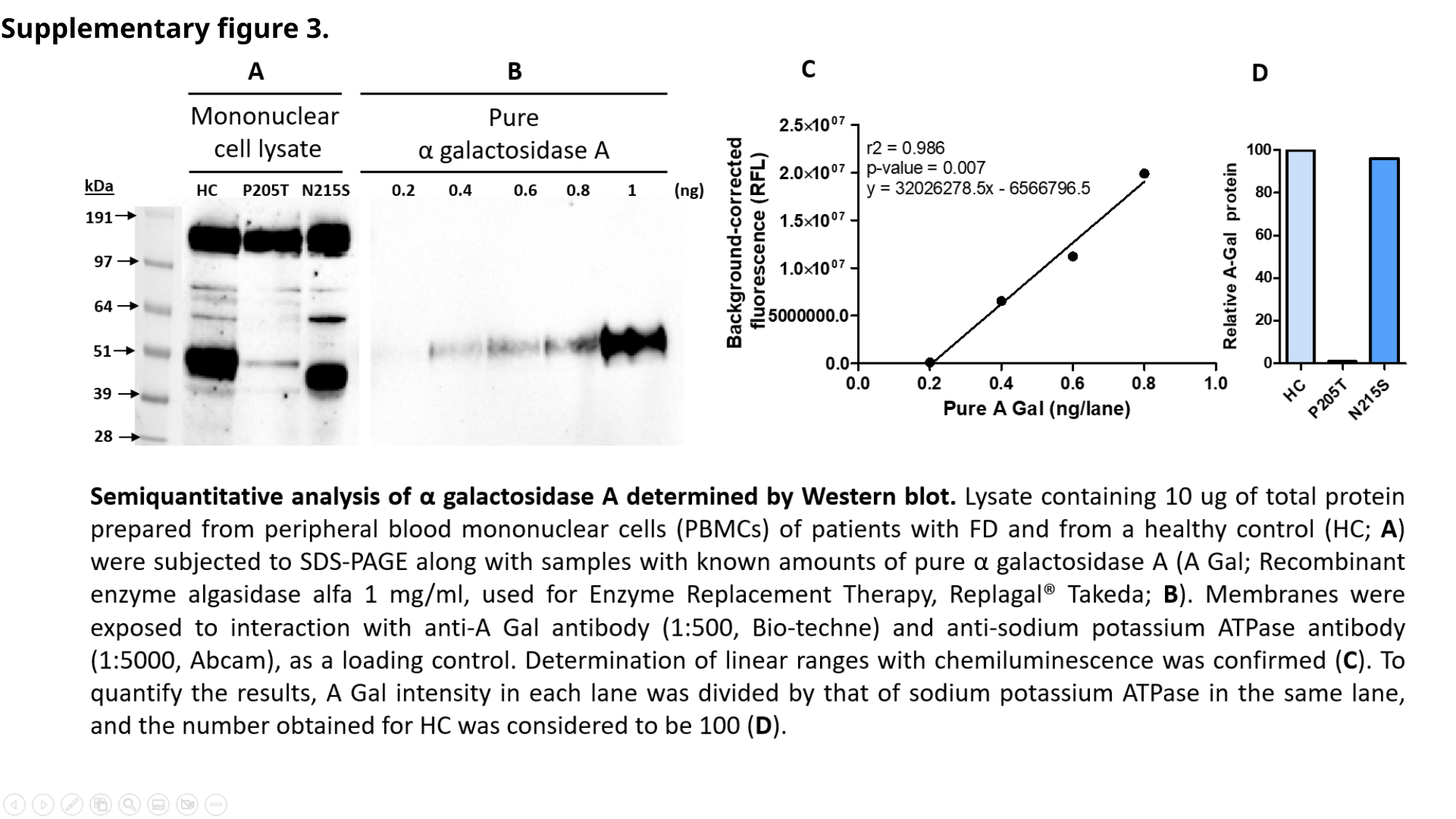

Supplementary figure 3.
#

### Slide 5
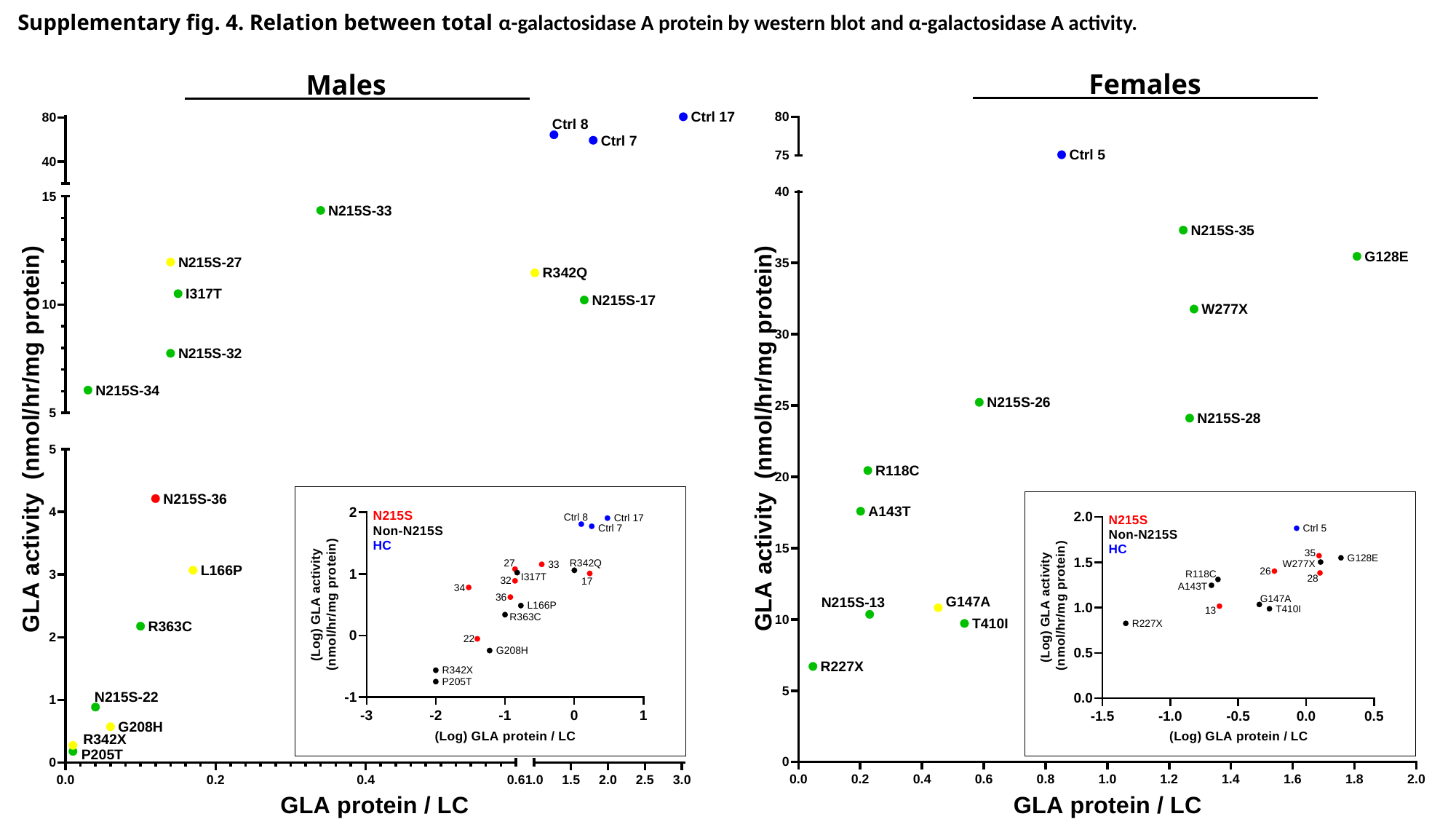

Supplementary fig. 4. Relation between total α-galactosidase A protein by western blot and α-galactosidase A activity.
Females
Males

### Slide 6
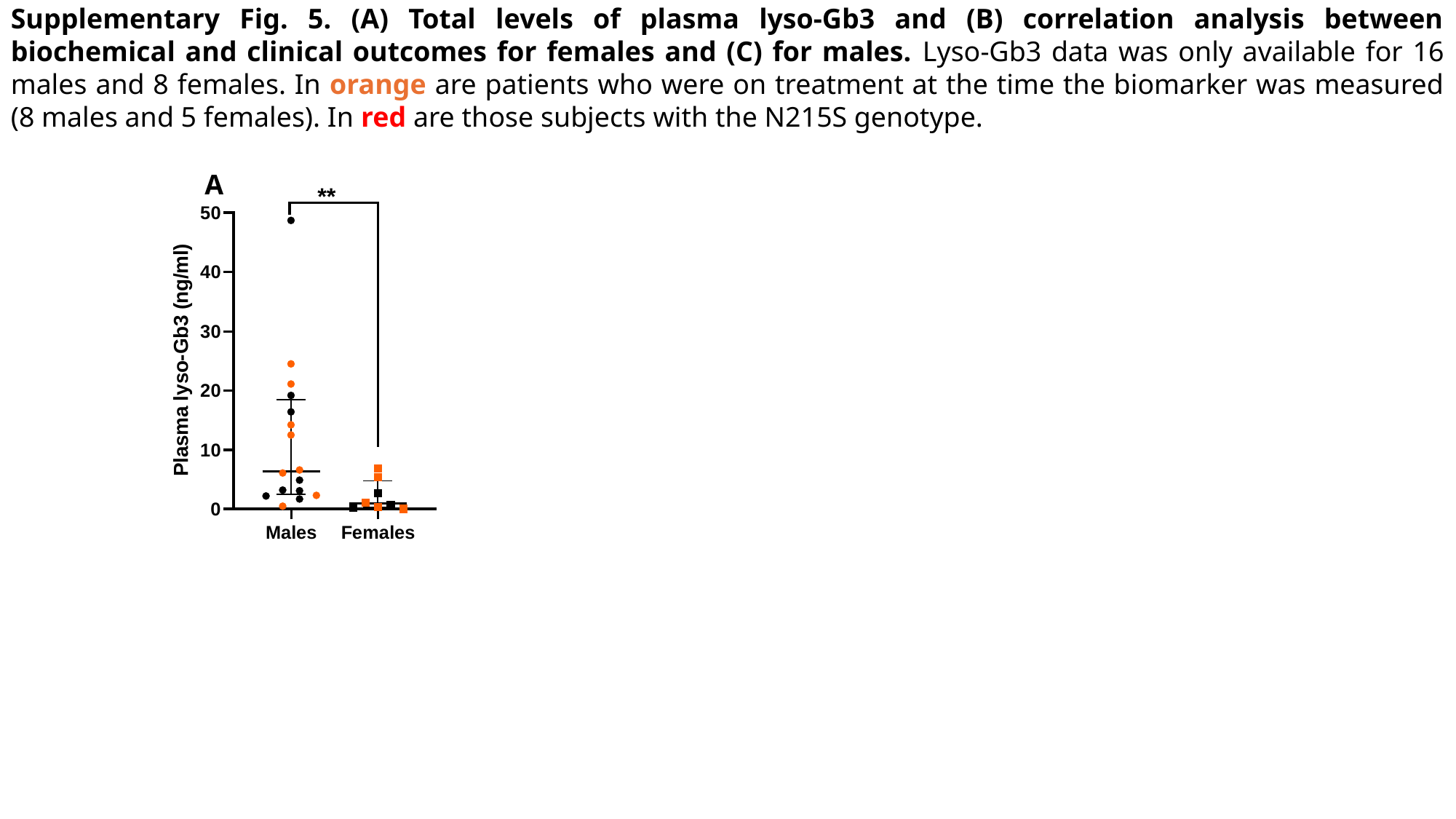

Supplementary Fig. 5. (A) Total levels of plasma lyso-Gb3 and (B) correlation analysis between biochemical and clinical outcomes for females and (C) for males. Lyso-Gb3 data was only available for 16 males and 8 females. In orange are patients who were on treatment at the time the biomarker was measured (8 males and 5 females). In red are those subjects with the N215S genotype.
A

### Slide 7
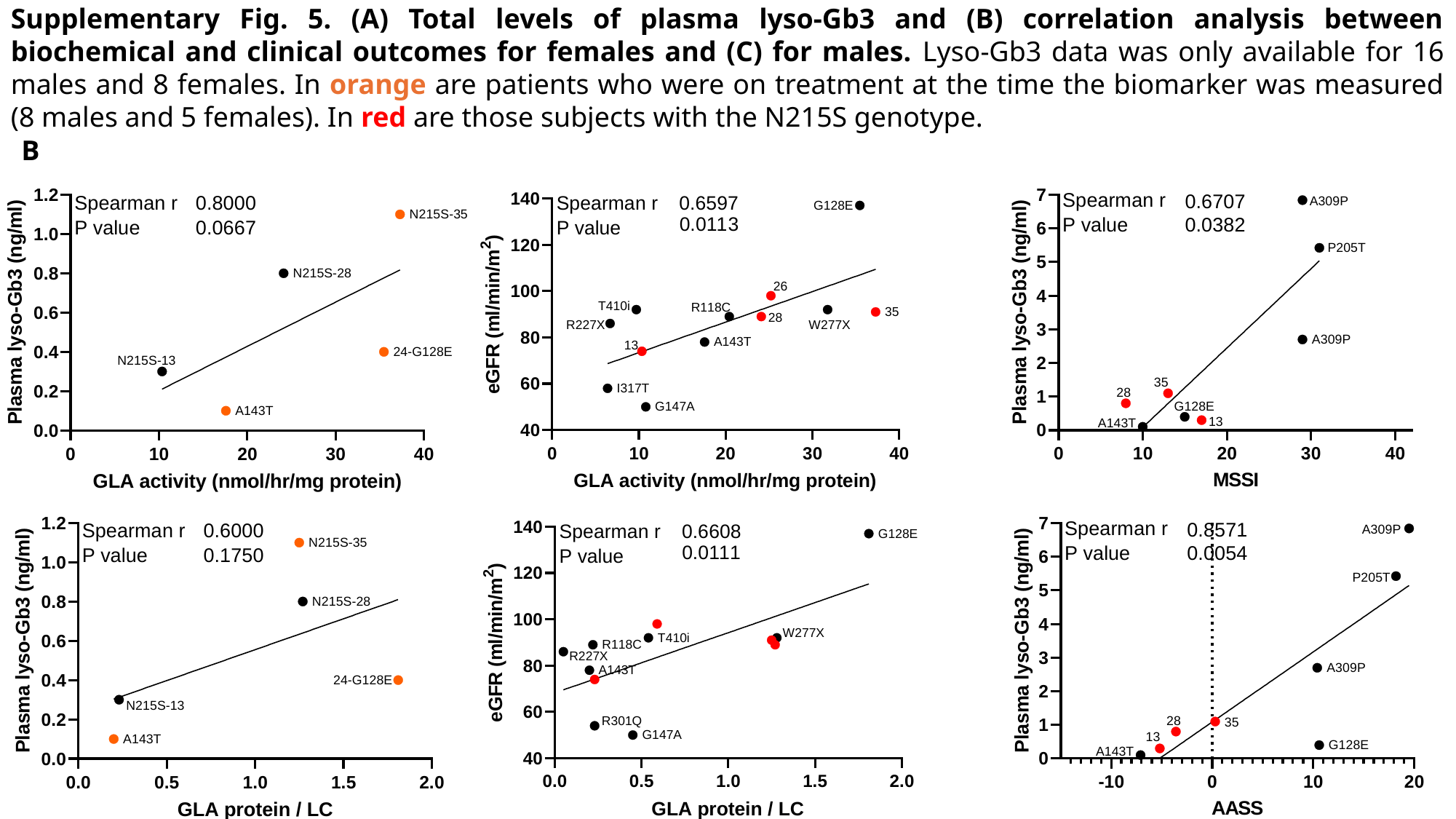

Supplementary Fig. 5. (A) Total levels of plasma lyso-Gb3 and (B) correlation analysis between biochemical and clinical outcomes for females and (C) for males. Lyso-Gb3 data was only available for 16 males and 8 females. In orange are patients who were on treatment at the time the biomarker was measured (8 males and 5 females). In red are those subjects with the N215S genotype.
B

### Slide 8
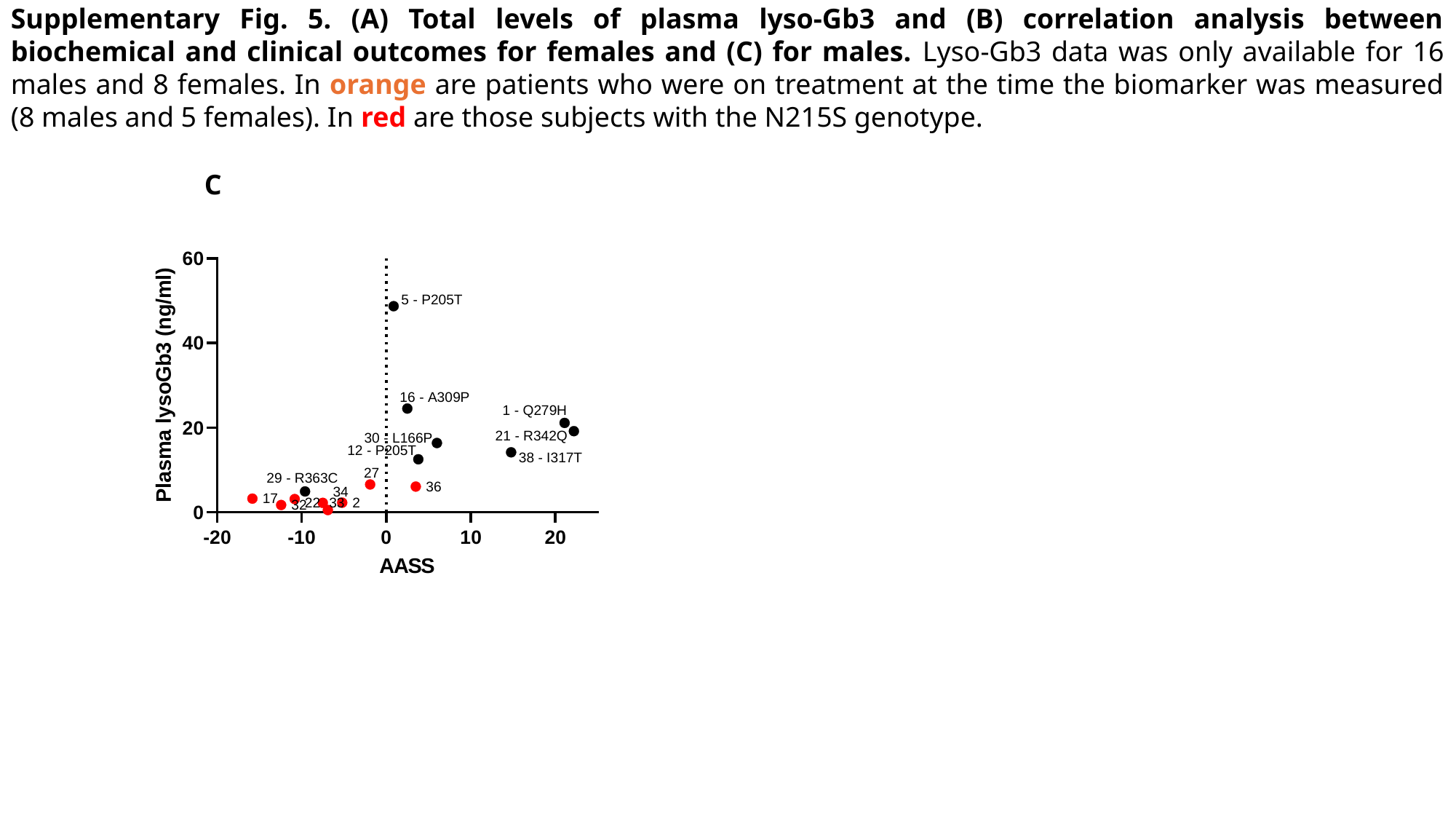

Supplementary Fig. 5. (A) Total levels of plasma lyso-Gb3 and (B) correlation analysis between biochemical and clinical outcomes for females and (C) for males. Lyso-Gb3 data was only available for 16 males and 8 females. In orange are patients who were on treatment at the time the biomarker was measured (8 males and 5 females). In red are those subjects with the N215S genotype.
C
